# Astrocytic FABP5 drives non-cell-autonomous oligodendrocyte injury in multiple system atrophy by promoting TNF signaling and ferroptotic stress

**DOI:** 10.1101/2025.11.08.687320

**Authors:** Chuantao Wu, Jiejing Lin, Yi Chen, Nozomi Takahashi, Qikai Chen, Baiyao Liu, Fangmeng Dai, Wenxue Zhao, David I Finkelstein, Ichiro Kawahata, Kohji Fukunaga, An Cheng

## Abstract

Multiple system atrophy (MSA) is a fatal α-synucleinopathy characterized by progressive parkinsonism, cerebellar and autonomic dysfunction. Currently, the mechanisms driving cerebellar white matter neuroinflammation and degeneration in MSA are poorly understood. Here, we identified fatty acid-binding protein 5 (FABP5) as a key factor regulating cerebellar inflammation in MSA pathogenesis in a detailed study of human, mouse and cultured astrocytes. Firstly, transcriptomic profiling of human MSA cerebellar white matter revealed activation of pro-inflammatory and ferroptotic pathways, with FABP5 identified as a key pathway component that is upregulated. We confirmed that FABP5 is upregulated in reactive astrocytes in the PLP-α-syn transgenic mouse model, also in LPS-treated primary astrocytes. Fabp5 silencing suppressed TNF signaling, mitigated ferroptosis, and restored mitochondrial function. These findings suggest astrocytic FABP5 as a central intracellular regulator linking glial inflammation, ferroptosis, and mitochondrial injury. Overall, this mechanism suggests that FABP5 drives pathology mediated by astrocytes oligodendroglia in MSA, therefore representing a novel and promising therapeutic target.

## 1. Introduction

Multiple system atrophy (MSA) is a rare, rapidly progressive, and fatal neurodegenerative disorder[1]. It is clinically characterized by a varying combination of severe autonomic failure, parkinsonism, and cerebellar ataxia[2]. Based on the dominant motor symptoms, MSA is classified into two major subtypes: the parkinsonian variant (MSA-P) and the cerebellar variant (MSA-C)[3]. The disease typically arises in the sixth decade of life, affects approximately 3-5 individuals per 100,000, and progresses relentlessly with a poor prognosis[4]. Pathologically, MSA belongs to the α-synucleinopathies but differs fundamentally from Parkinson’s disease (PD) and dementia with Lewy bodies (DLB), as misfolded α-synuclein (α-syn) accumulates primarily within oligodendrocytes, forming glial cytoplasmic inclusions (GCIs)[5–8]. A critical feature of MSA is the dysfunction and loss of oligodendrocytes, leading to impaired myelin maintenance and subsequent oligodendrocyte white-matter degeneration[9–11]. Oligodendrocyte degeneration in MSA is thought to arises from a combination of α-syn toxicity, mitochondrial dysfunction, oxidative stress, and lipid dysregulation[12]. In parallel, α-syn transferred from neurons is thought to accumulates in oligodendrocytes and forms GCIs, initiating oligodendroglial degeneration[12]. Lipid metabolic imbalance further increases cellular vulnerability by promoting membrane instability and oxidative injury.

Recent studies suggest that the propagation of misfolded α-syn plays a central role in oligodendrocyte degeneration in MSA. Neuron-derived α-syn can be internalized by oligodendrocytes, where it accumulates to form toxic aggregates and contributes to the development of GCIs[13]. Emerging evidence indicates that FABP7 facilitates the formation and dissemination of cytotoxic α-syn hetero-aggregates through clathrin-dependent endocytosis, thereby exacerbating oligodendroglial loss and cerebellar dysfunction[14]. Moreover, focal neuroinflammation within cerebellar white matter has been reported in ataxic conditions and is sufficient to induce motor impairment, consistent with observations in acute cerebellitis-associated ataxia[15]. In MSA, diffusion magnetic resonance imaging (MRI) and microstructural imaging studies further reveal widespread impairment of cerebellar-related white-matter pathways, and alterations within these tracts correlate with the severity and progression of motor deficits in affected individuals[16, 17].

Beyond MSA, neuroinflammation is a shared pathological feature across multiple neurodegenerative diseases, where chronic activation of microglia and astrocytes drives oxidative stress and neuronal damage, often through the secretion of pro-inflammatory cytokines such as IL-1β, TNF-α, and IL-6[18–21]. Crosstalk between reactive glial populations amplifies lipid peroxidation and compromises oligodendroglia viability. Despite being well characterized in PD and DLB, glial-driven inflammation in MSA remains less explored[12]. Given that oligodendrocyte loss and demyelination are central pathological features of MSA, inflammatory responses are likely not merely secondary consequences but active contributors to disease progression. This gap highlights an unmet need to understand how glial metabolic regulators orchestrate inflammatory and degenerative signaling within cerebellar oligodendrocyte and white matter in MSA.

Fatty acid-binding proteins (FABPs) are intracellular lipid chaperones that regulate the trafficking and partitioning of long-chain fatty acids, thereby influencing membrane composition, redox vulnerability, and inflammatory signaling[22]. Among them, FABP5 has been increasingly implicated in the integration of lipid metabolism and oxidative inflammatory responses in the central nervous system. FABP5 enriches cellular membranes with oxidation-prone polyunsaturated fatty acids (PUFAs), sensitizing glial cells to lipid peroxidation and ferroptotic stress[23]. In oligodendrocytes, FABP5 interacts with VDAC-1 and BAX to form mitochondrial macropores, leading to cytochrome-c and mtDNA release and increased IL-1β under psychosine stress[24]. Mitochondrial accumulation of FABP5 also promotes lipid peroxidation and BAX/VDAC-1 channel formation in neurons, aggravating oxidative injury[25]. In parallel, FABP5 activity enhances ferroptosis susceptibility under hypoxic and iron-rich conditions by facilitating PUFAs incorporation into phospholipid[26]. These findings indicate that FABP5 integrates lipid transport, oxidative lipid stress, inflammatory cascades, and ferroptotic vulnerability, making it a compelling candidate for mechanistic investigation within the cerebellar white matter of MSA.

In the present study, we demonstrate that astrocytic FABP5 acts as a key regulator involved in cerebellar white matter dysfunction, which contribute to the specific glial MSA pathogenesis. Our transcriptomic and histological analyses of MSA donor white matter and PLP-αSyn mice cerebellum reveals that FABP5 is involved in pro-inflammatory, anti-survival signaling and ferroptosis pathway in cerebellar oligodendrocyte and white matter of MSA. Furthermore, FABP5 silencing mitigates FABP5-mediated astrocytic inflammation, ferroptosis, and even mitochondrial injury, highlighting the potential of FABP5 to be a novel therapeutic approach for MSA.

## 2. Materials and methods

### 2.1 Animal

Pregnant C57BL/6J mice were obtained from the Laboratory Animal Center of Sun Yat-sen University (Shenzhen, China) and housed in polypropylene cages under specific pathogen-free (SPF) conditions (temperature: 23 ± 2 ℃, humidity: 55 ± 5%, lights on from 9 a.m. to 9 p.m.). Animals had unlimited access to food and water. All animal experiments were reviewed and approved by the Institutional Animal Care and Use Committee of Sun Yat-sen University (SYSU-IACUC-2025-001716).

Male and female PLP-α-syn mice overexpressing human wild-type α-syn under the control of the oligodendrocyte-specific proteolipid protein promoter (PLP-α-syn mice) on a C57Bl/6 background were used for all experiments. The mice were bred housed in a temperature-controlled, pathogen-free environment under a 12-h light/dark cycle at the animal facility of The Florey Institute of Neuroscience and Mental Health and allowed access to standard laboratory chow and water ad libitum. All the experiments were approved by the Florey Institute Animal Ethics Committee (13-117 and #18-006)[27, 28].

All animal procedures were conducted in accordance with the guidelines of the Florey Institute of Neuroscience and Mental Health Animal Ethics Committee and the Australian Code for the Care and Use of Animals for Scientific Purposes. Experimental protocols were approved under the following ethics approvals. For euthanasia, the mice were intraperitoneally injected with an overdose of sodium pentobarbital. All efforts were made to minimize animal suffering and minimize the number of animals used.

### 2.2 Primary astrocyte isolation

Mouse astrocytes were isolated and cultured as previously described [29]. Briefly, P1-2 mouse pups were decapitated and cortex from the brains was dissected and immersed in PBS containing 0.5mM EDTA. The cortices then were diced using a sterilized razor blade into ∼1 mm^3^ chunks and digested in 0.05% trypsin for 15 min at 37℃. The tissue suspension then was passed through a 70 μm nylon cell strainer. Cells were centrifuged (300 g, 5 min) and resuspended in Dulbecco’s Modified Eagle’s Medium (DMEM, 4 mM L-glutamine, 20% FBS, 50 U mL-1 penicillin, and 50 μg/mL streptomycin), seeded in tissue culture T25 flasks and cultured in DMEM medium. After 10 days, the flasks were shaken for 18 h at 180 rpm at 37℃ to remove microglial cells and oligodendrocyte, and the residual cells were considered crude astrocyte cultures. Finally, astrocytes were stained with GFAP. Ethical approval was obtained from the Institutional Animal Care and Use Committee of Sun Yat-sen University (SYSU-IACUC-2025-001716).

### 2.3 FABP5 shRNA plasmid and transfection

Three mouse FABP5-targeting shRNA expression plasmids (pRNAi-U6neo bcakbone) were obtained from Corues Biotechnology (Nanjing, China). The target sequence was as follows: shRNA1: GATGGAAAGCCACGGCTTTGACTCGAGTCAAAGCCGTGGCTTTCCATCTTTTTT; shRNA2: GGGAAGGAGAGCACGATAACACTCGAGTGTTATCGTGCTCTCCTTCCCTTTTTT; shRNA3: GGAGTGTGTCATGAACAATGCCTCGAGGCATTGTTCATGACACACTCCTTTTTT. Plasmid were amplified in *E.coli* DH5α and purified using an Endo-Free Plasmid DNA Maxi Kit (OMEGA) according to the manufacturer’s instructions. Primary astrocytes were seeded at a density of 5 × 10^5 cells per dish into 35mm dishes 48 h prior to transfection to achieve ∼70–90% confluence on the day of transfection. Cells were transfected using Lipofectamine LTX and PLUS Reagent (Invitrogen) with Opti-MEM (Gibco) according to the manufacturer’s protocol. Briefly, 2 μg plasmid DNA was mixed with 2 μL PLUS reagent in 700 μL Opti-MEM and incubated for 5 min at room temperature. Separately, 2 μL LTX reagent was diluted in 100 μL Opti-MEM. The two solutions were combined and incubated for 15 min to allow complex formation, then added to cells. After 6 hours, the transfection medium was replaced with complete astrocyte culture medium.

### 2.4 Protein extraction and western blotting (WB)

Cells were washed twice with ice-cold PBS and then directly scraped into RIPA lysis buffer (Beyotime) supplemented with protease (APExBIO) and phosphatase inhibitors (APExBIO). The collected lysates were transferred to pre-chilled 1.5 mL tubes and ultrasonicated in an ice-water bath for 10 min at 4 °C. The lysates were then centrifuged at 15,000 g for 10 min at 4 °C and the supernatant was collected.

The protein samples that were prepared for WB were mixed with 5 × SDS loading buffer and incubated for 10 min at 100 °C. Equal sample volumes (10 μL per lane) were loaded and separated, using SDS-polyacrylamide gel electrophoresis and SDS–polyacrylamide gels spanning 12.5-15% (EpiZyme Biotechnology). The gels were then transferred to PVDF membranes (0.45 μm, Millipore, Cat. No. IPVH00010) in a buffer containing 3.03 g/L Tris, 14.41 g/L glycine using a wet transfer system (100 V, 60 min). The membranes were blocked in 5% defatted milk in TBST solution containing 10 mM Tris–HCl (pH 7.5), 150 mM NaCl, and 0.1% Tween-20 for 1 h at room temperature. Membranes were incubated overnight at 4°C with primary antibodies diluted in TBST: mouse anti-FABP5 (1:1000, Proteintech), rabbit anti-GPX3 (1:1000, Proteintech), rabbit anti-TNF-α (1:1000, Proteintech), rabbit anti-ACSL5 (1:1000, Proteintech), rabbit and β-Actin (1:50000, Huabio). After washing three times with TBST (10 min each time), membranes were incubated with TBST-diluted appropriate HRP-conjugated secondary antibodies for 1 h at room temperature. Bands were visualized using an ECL (Enhanced Chemiluminescence) luminescence detector (Absin) and quantified by ImageJ. Protein expression levels were normalized to β-Actin.

### 2.5 Immunofluorescent (IF) staining

Purified astrocyte cells were cultured on tissue culture glass slides in 12-well dishes and fixed in 4% paraformaldehyde for 30 min at room temperature. Glass slides were washed three times with PBS/1% BSA (10 min each time), and permeabilized with 0.1% Triton X-100 in PBS for 10 min, then blocked in PBS/1% BSA for 1 h at 4 °C. Glass slides were incubated with primary antibody against GFAP (1:500) in a blocking solution at 4°C overnight. After washing three times with PBS/1% BSA (10 min each time), glass slides were incubated with Alexa 568-labeled anti-mouse IgG H&L secondary antibody (1:1000) overnight at 4°C in the dark. Then glass slides were washed three times with PBS/1% BSA (10 min each time), and counterstained with DAPI for 15 min at room temperature to stain the nucleus. Finally, glass slides were mounted with the fluoroshield mounting medium. The astrocyte cells were observed and photographed with a confocal laser scanning microscope (DMi8; Leica, Wetzlar, Germany).

### 2.6 RNA sequencing and analysis

Cultured cells were digested in 0.25% trypsin and collected into RNase-free microcentrifuge tubes and kept on ice. Cells then were centrifuged (300 g, 5 min, 4℃) and resuspended in sterile PBS, and centrifuged (300 g, 5 min, 4℃) again. The supernatants were removed completely, and the microcentrifuge tubes with cell pellet were placed in liquid nitrogen for 15 to 30s. Samples were submitted to Novogene (Beijing, China) for bulk RNA sequencing using Illumina NovaSeq6000 platform.

### 2.7 Bioinformatics and data visualization

RNA-seq data of MSA cerebellar white matter (n=47 MSA, n=47 controls; GEO: GSE199715) [30] and laser-captured human oligodendrocytes (GSE199724)[30] were obtained from GEO. Bulk RNA-seq counts from GSE199715 were initially analyzed using the GEO2R platform for differential expression. 1 Adjusted p-values (Benjamini–Hochberg FDR), log2 fold changes, and p-values were automatically generated. 1 For both this public dataset and the in-vitro astrocyte RNA-seq data, Gene Ontology (GO) and KEGG pathway enrichment analyses of differentially expressed genes (DEGs, defined as adjusted p-value < 0.05) were performed using the DAVID Functional Annotation Tool.

To delineate transcriptomic modules in the GSE199715 dataset, Weighted Gene Co-expression Network Analysis (WGCNA) was performed (Fig. S1). Genes were clustered using hierarchical clustering based on topological overlap, and co-expression modules were identified using the dynamic tree cut method; module eigengenes (MEs) were then correlated with MSA disease status using Pearson correlation.

A separate FABP5-centered co-expression analysis was performed using Pearson correlation (|r| > 0.2, FDR < 0.05). For cell-type deconvolution of GSE199715, a reference signature matrix was generated from scRNA-seq data (GEO: GSE67835) [], including astrocytes (n=49), microglia (n=9), and oligodendrocytes (n=38). Both datasets were normalized to Transcripts Per Million (TPM). The Non-negative least squares (NNLS) algorithm in R was applied to estimate cell-type proportions, and group differences were evaluated using unpaired two-tailed t-tests. For visualization of gene expression in mouse brain (Fig. 3D), t-SNE plot data was sourced from SRA850958:SRS4386089. Protein-protein interaction network analysis (Fig. 4F) was conducted using the STRING database.

All visualizations, including variance-stabilizing transformation (VST) for normalization, volcano plots, Principal Component Analysis (PCA), hierarchical clustering heatmaps, scatter plots, and correlation matrices, were performed in R (v4.5.0) using packages including ggplot2 and heatmaps.

### 2.8 Lipid ROS assay

Lipid ROS levels in primary astrocytes were assessed using the BODIPY 581/591 C11 probe (Beyotime). Cells were divided into three groups: Control, LPS (1 μg/mL, 24 h), and LPS + FABP5 silencing (FABP5 shRNA transfection for 5 days followed by LPS exposure). Additional controls included unstained samples for fluorescence gating. On Day-1, astrocytes were equally seeded into a 24-well plate. On Day 0, FABP5 shRNA plasmids were transfected into the designated group. On Day 4, LPS was added to induce an inflammatory environment. On Day 5, cells were harvested by trypsinization, centrifuged at 300 × g for 5 min at 4°C, and washed twice with cold PBS. Cell pellets were incubated with 5 μM BODIPY 581/591 C11 diluted in PBS for 30 min at 37°C in the dark. After incubation, samples were washed twice at 4°C and resuspended in ice-cold PBS.

Flow cytometry was performed using Beckman Coulter CytoFLEX platform equipped with 488-nm laser. The oxidized (510 nm) signals were detected through FITC channel. Data were analyzed using FlowJo software. Lipid ROS levels were quantified as mean fluorescence intensity (MFI) and statistically evaluated by one-way analysis of variance (ANOVA).

### 2.9 Statistics

Data were analyzed using GraphPad Prism 10 (GraphPad Software, Inc., La Jolla, CA, USA) and are expressed as mean ± standard error of the mean (SEM). The normality of the data was assessed using the Shapiro-Wilk test, and homogeneity of variances was evaluated using the Brown-Forsythe test, both in GraphPad Prism 10. For data meeting these assumptions, significant differences were determined using Student’s t-test for two-group comparisons and one-way or two-way ANOVA for multigroup comparisons, followed by Tukey’s multiple comparisons test. Statistical significance was set at p < 0.05. For randomization and blinding, we used double-blind trials. Briefly, the mice were randomly assigned to control or experimental groups and cultured or treated by Experimenter A. During the imaging analysis, other experimenters did not know the real group information.

## 3. Results

### 3.1 A pro-inflammatory transcriptomic signature in MSA cerebellar white matter identifies FABP5 as a key upregulated node

To identify the key molecular drivers of the cerebellar white matter oligodendrocyte pathology observed in MSA, our investigation began at the transcriptomic level. To characterize transcriptional alterations associated with MSA, we analyzed publicly available RNA-seq data (GSE199715) from cerebellar white matter of MSA patients (n=47) and healthy controls (n=47) (Fig. 1A). KEGG enrichment of differentially expressed genes (DEGs) revealed a selective activation of inflammatory signaling, with TNF and VEGF pathways most prominently upregulated in MSA samples (Fig. 1B), consistent with previously identified inflammatory signatures in MSA cerebellar white matter [31].

**Figure 1.**
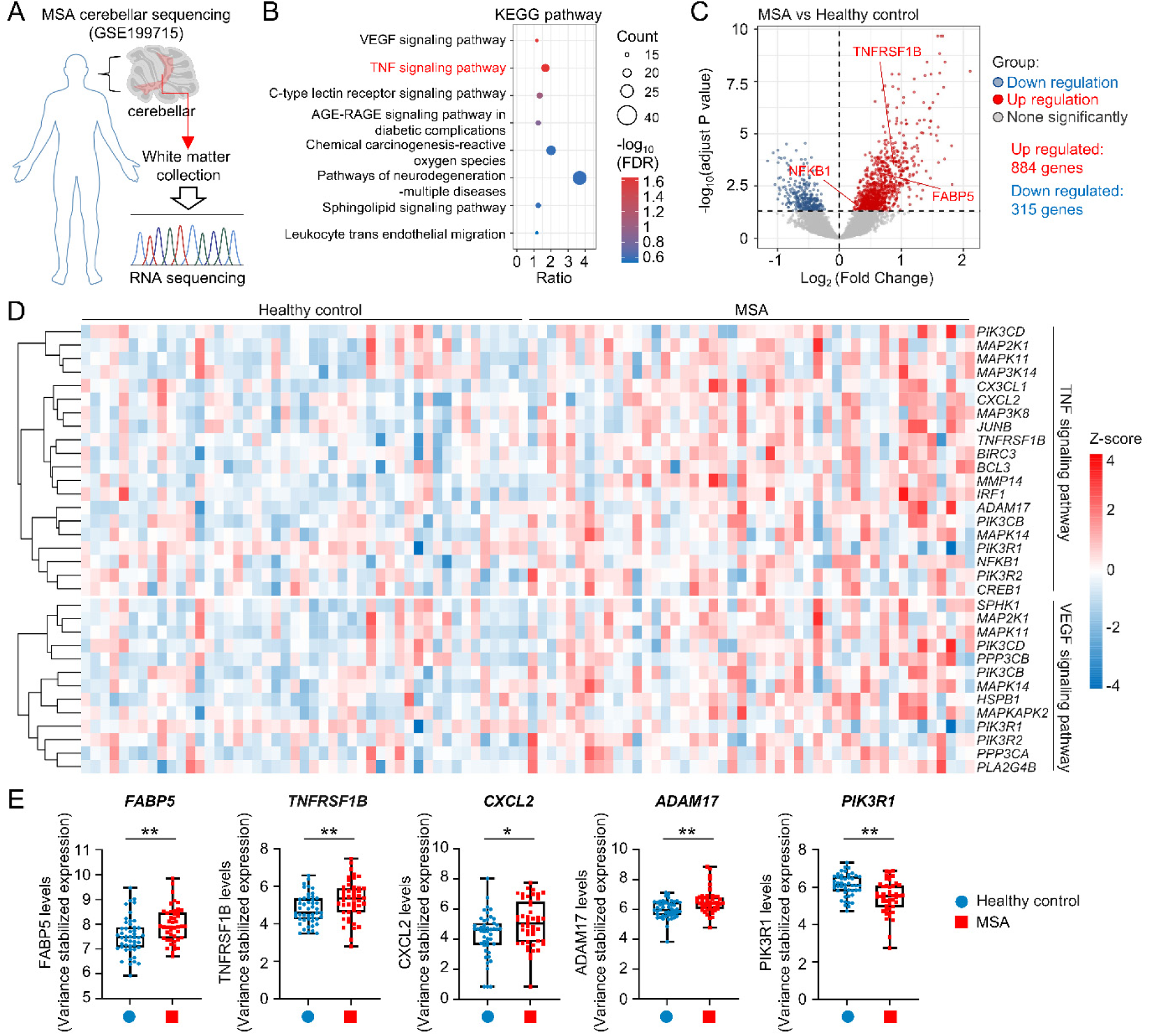
Transcriptomic profiling reveals a pro-inflammatory and anti-survival signature in the cerebellar white matter of MSA. (A) Schematic overview of the study design. Cerebellar white matter tissue from multiple system atrophy (MSA) patients and healthy controls was collected and subjected to RNA sequencing (dataset GSE199715). (B) KEGG pathway enrichment analysis of differentially expressed genes (DEGs) between MSA and healthy controls. Pathways significantly enriched (FDR < 0.05) are shown, with color indicating false discovery rate (FDR) and dot size indicating the number of genes involved. Notably, VEGF and TNF signaling pathways were significantly upregulated in MSA. (C) Volcano plot showing DEGs in MSA versus healthy controls. Red dots represent significantly upregulated genes, blue dots indicate significantly downregulated genes, and gray dots represent genes not significantly altered. Key upregulated genes related to TNF signaling pathways are labeled (e.g., *FABP5*, *TNFRSF1B*). A total of 884 genes were upregulated and 315 were downregulated (adjusted p-value < 0.05). (D) Heatmap showing hierarchical clustering of representative DEGs involved in VEGF and TNF signaling pathways. Expression levels (Z-scores) are visualized across samples from healthy controls and MSA patients. (E) Boxplots showing variance stabilized expression levels of selected representative DEGs: *FABP5*, *TNFRSF1B*, *CXCL2*, and *ADAM17* with significant upregulation and *PIK3R1* with significant downregulation in MSA samples compared to controls. Data are presented as boxplots (showing median, interquartile range, and individual data points), with Healthy controls (n=47) and MSA patients (n=47). Significance stars represent the adjusted p-value (padj) from DESeq2. *p < 0.05, **p < 0.01.

Differential expression analysis revealed 884 upregulated and 315 downregulated genes in MSA versus controls (Fig. 1C). Among the significantly upregulated genes, *FABP5* showed a notable increase, together with several TNF-related genes such as *TNFRSF1B* and *NFKB1*, indicating a pronounced inflammatory transcriptional shift. Hierarchical clustering of representative TNF and VEGF pathway genes clearly further distinguished MSA and healthy controls, reflecting a transcriptomic divergence between two groups (Fig. 1D). Moreover, variance-stabilized expression analysis confirmed elevated levels of *FABP5*, *TNFRSF1B*, *CXCL2*, and *ADAM17*, alongside reduced expression of *PIK3R1* in MSA (Fig. 1E). Given previous evidence linking FABP5 to glial inflammatory activation and oxidative injury [32, 33], its selective upregulation positions it as a potential contributor to neuroinflammatory pathology in MSA.

Taken together, these integrated transcriptomic analyses delineate a pro-inflammatory and anti-survival gene expression program in the cerebellar white matter of MSA and identify FABP5 as a distinctly upregulated gene that may contribute to glial inflammatory activation in the disease.

### 3.2 The FABP5 co-expression network functionally links inflammation and ferroptosis in MSA

Having identified *FABP5* as a key upregulated gene in the overall MSA inflammatory signature (Fig. 1E), we next sought to define its specific functional network. To delineate transcriptomic modules altered in MSA cerebellar white matter, we first performed weighted gene co-expression network analysis (WGCNA) on GSE199715 (Fig. S1). The gene dendrogram and color-coded module map (Fig. S1A) showed multiple co-expression modules derived from hierarchical clustering. The module-trait relationship heatmap (Fig. S1B) highlighted that several modules exhibited strong correlations with MSA, with the pink and black modules showing the most significant negative correlations, implying downregulation of genes related to cellular homeostasis. Scatter plots of module membership versus gene significance (Fig. S1C) confirmed a coherent internal structure within these modules, validating the reliability of the network.

Based on the disease-associated modules identified by WGCNA, we next explored genes showing expression patterns closely related to MSA pathology. Because FABP5 was significantly upregulated in MSA (Fig. 1E), we next characterized its co-expression context. FABP5 was positively co-expressed with 6,885 genes (Fig. 2A), and 352 of these overlapped with DEGs between MSA and control samples (Fig. 2B). Crucially, this analysis provided our first major mechanistic insight: KEGG analysis of the overlapping genes highlighted pathways related to TNF signaling pathway and ferroptosis, implying that FABP5 is embedded in a molecular program coupling inflammatory activation and iron-dependent lipid peroxidation.

**Figure 2.**
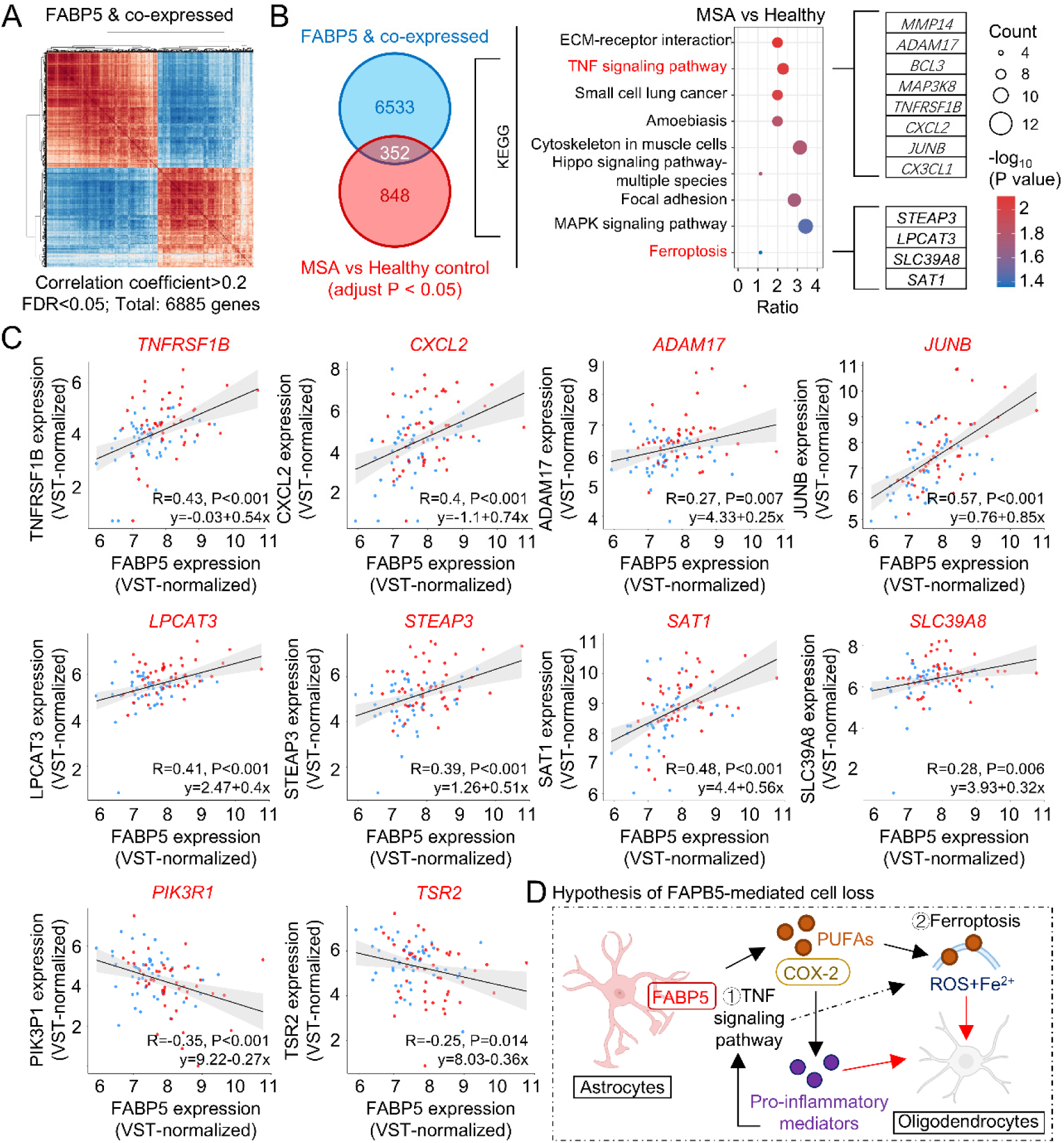
Co-expression analysis of FABP5 in MSA reveals links to inflammation and ferroptosis, and a proposed mechanism for FABP5-mediated cell loss. (A) Pearson correlation heatmap showing genes co-expressed with FABP5 (a total of 6,885 genes) based on the criteria: |correlation coefficient| > 0.2, FDR < 0.05. (B) Left, Venn diagram illustrates the overlap between FABP5 co-expressed genes and differentially expressed genes in multiple system atrophy (MSA) patients compared to healthy controls (adjusted P < 0.05), identifying 352 shared genes. Right, KEGG pathway enrichment analysis of the overlapping genes, highlighting significant enrichment in pathways such as TNF signaling and ferroptosis. Dot size indicates the number of genes per pathway, and color represents statistical significance (-log10 adjusted P-value). (C) Scatter plots showing the correlation between FABP5 expression (VST-normalized) and expression of key representative genes from the enriched pathways, including TNF signaling pathway (TNFRSF1B, CXXCL2, ADAM17, JUNB), Ferroptosis pathway (LPACT3, STEAP3, SAT1, SLC39A8) and cell survival related genes (PIK3R1, TSR2). Pearson correlation coefficients (R) and p-values are shown on each plot. (D) Hypothesis of FABP5-mediated oligodendrocyte cell loss: In astrocytes promotes the metabolism of polyunsaturated fatty acids (PUFAs) and activates the TNF/COX-2 signaling pathway, leading to increased release of pro-inflammatory mediators and induction of ferroptosis, thereby contributing to oligodendrocyte injury.

**Figure 3:**
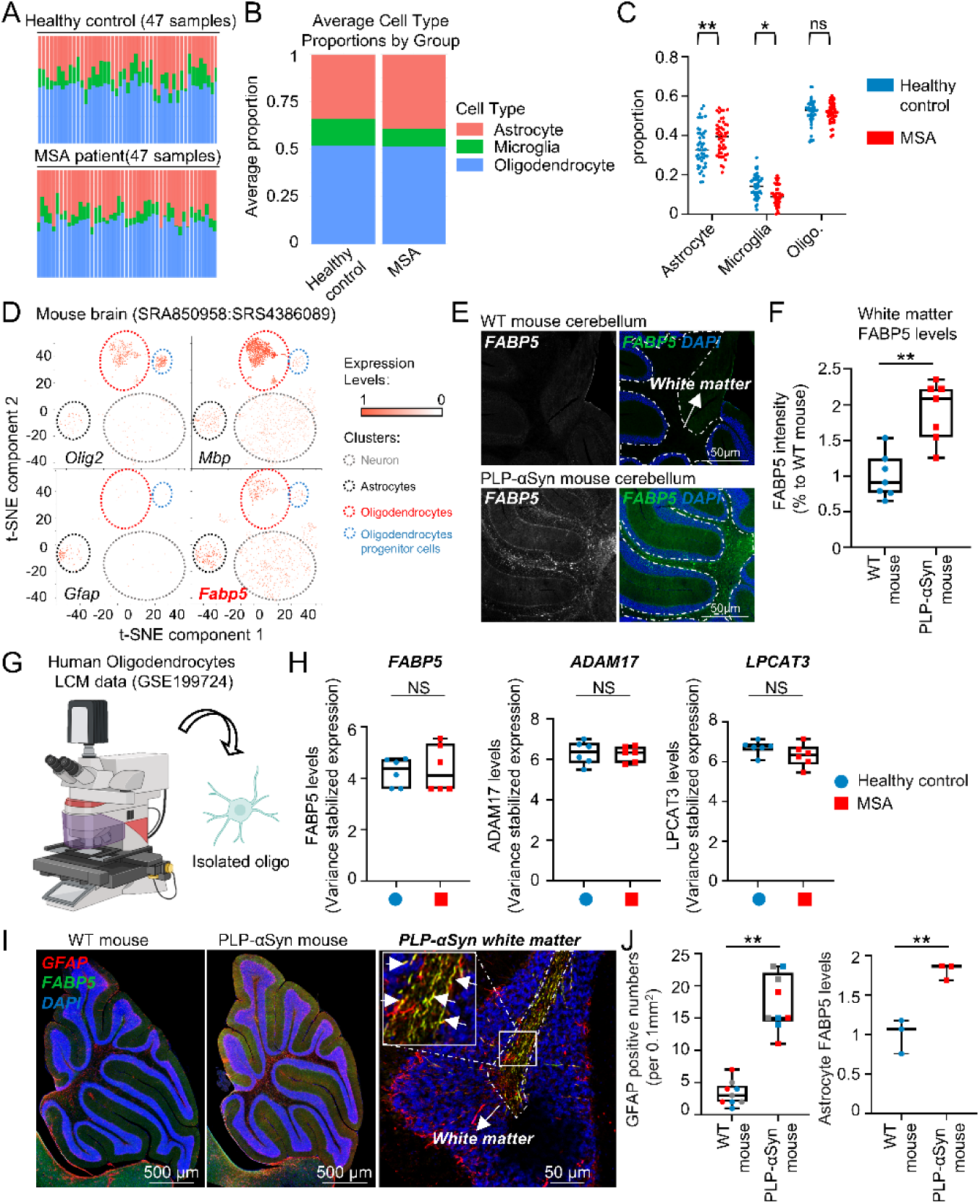
Altered glial cell composition and FABP5 expression in MSA patients and mouse models. (A) Bar plots showing the proportion of astrocytes (red), microglia (green), and oligodendrocytes (blue) across 47 healthy controls (top) and 47 MSA patient cerebellum white matter samples (bottom). (B) Average cell type proportions for each group, revealing an increase in astrocytes, but decrease in microglia in MSA white matter compared to controls. (C) Box plots quantifying the differences in cell type proportions between healthy controls (blue) and MSA patients (red). Statistical significance was determined using multiple unpaired t-tests (*p < 0.05, **p < 0.01). (D) t-SNE plots of single-cell RNA-seq data from mouse brains, displaying gene expression of oligodendrocyte markers (*Olig2*, *Mbp*), astrocyte marker (*Gfap*), and *Fabp5*. *Fabp5* expression is enriched in oligodendrocyte and astrocyte clusters. (E) Representative immunofluorescence images showing Fabp5 expression (white) and DAPI (blue) in cerebellar white matter from wild-type (WT) and PLP-α-syn mice. Elevated Fabp5 levels are observed in PLP-α-syn mouse white matter. (F) Quantification of Fabp5 fluorescence intensity in cerebellar white matter. Data are presented as boxplots; **p < 0.01 by unpaired t-test. (G) Schematic of the experimental workflow: Isolation of oligodendrocytes from human samples (GSE199724) via Laser Capture Microdissection (LCM). (H) Comparison of *FABP5*, *ADAM17*, and *LPCAT3* gene expression levels (variance stabilized expression) in human oligodendrocytes isolated in (G). NS (Not Significant) indicates no significant difference (by unpaired t-test). (I) Multiplex immunofluorescence staining for GFAP (red, astrocytes), FABP5 (green), and DAPI (blue) in the cerebellum of WT mice and PLP-α-syn mice. (J) Box plots quantifying the GFAP positive cell (astrocyte) numbers and FABP5 levels in astrocytes, showing increased FABP5 expression in astrocytes in PLP-α-syn mice. Data are presented as boxplots; **p < 0.01 by unpaired t-test.

**Figure 4.**
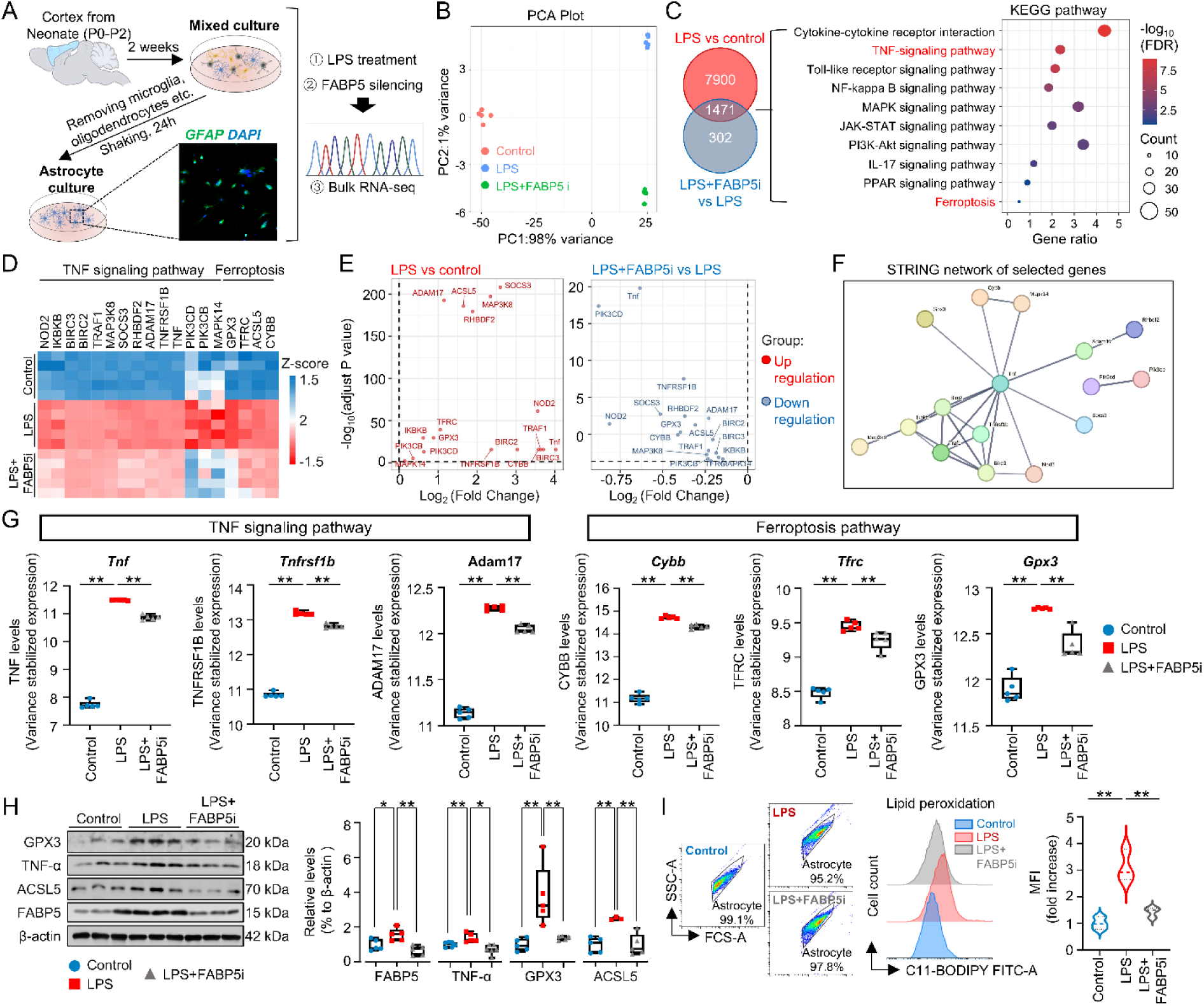
FABP5 silencing alleviates LPS-induced inflammation and ferroptosis in astrocytes. (A) Schematic diagram of the experimental workflow. Primary astrocytes were isolated from the cortices of neonatal (P0-P2) mice, purified, and cultured. Astrocyte identity was confirmed by immunofluorescence staining (GFAP in green; DAPI for nuclei in blue). The cultured astrocytes were divided into three groups for subsequent experiments: (1) Control, (2) Lipopolysaccharide (LPS) treatment, and (3) LPS treatment combined with FABP5 silencing. Bulk RNA-sequencing (RNA-seq) was then performed. (B) Principal Component Analysis (PCA) plot based on the RNA-seq data from the three experimental groups (Control, LPS, LPS+FABP5 silencing). The results show clear separation between the groups along Principal Component 1 (PC1, 98% variance) and Principal Component 2 (PC2, 1% variance). (C) Analysis of differentially expressed genes (DEGs). The Venn diagram (left) shows the number of unique and overlapping DEGs between the “LPS vs control” and “LPS+FABP5i vs LPS” comparisons. The bubble plot (right) displays the KEGG pathway enrichment analysis for the 1471 overlapping DEGs, revealing significant enrichment in the TNF signaling pathway and the Ferroptosis pathway. Bubble size represents the gene count, and the color scale indicates the enrichment significance (-log10(FDR)). (D) Heatmap showing the Z-score normalized expression levels of key genes from the TNF signaling and ferroptosis pathways across the three experimental groups. Red indicates upregulation, and blue indicates downregulation. The results demonstrate that LPS induces the expression of these genes, while FABP5 silencing reverses this effect. (E) Volcano plots visualizing the DEGs for the “LPS vs control” (left panel) and “LPS+FABP5i vs LPS” (right panel) comparisons. The plots show that many pro-inflammatory and ferroptosis-related genes (e.g., Tnf, Socs3, Tfrc) upregulated by LPS are significantly downregulated upon FABP5 silencing. (F) STRING protein-protein interaction network analysis of key genes from the TNF signaling and ferroptosis pathways, showing their functional relationships. (G) Plots of variance-stabilized expression levels for key TNF signaling genes *(Tnf, Tnfrsfl b,* and *Adami* 7) and ferroptosis genes *(Cybb, Tfrc,* and *Gpx3)* from the RNA-seq data. Compared to the control group, LPS significantly increased the expression of selected genes. FABP5 silencing significantly inhibited this LPS-induced upregulation. Data are presented as mean ±SEM (for bar charts) or as box plots (for *Gpx3).* Significance stars represent the adjusted p-value (padj) from DESeq2. **p < 0.01. (H) Western blot and quantification of FABP5, GPX3, ACSL5 and TNF-α protein levels in primary astrocyte, suggested FABP5 silencing (LPS+FABP5i) significantly reduced TNF-α expression and ACSL5, GPX3 levels, induced by LPS. (I) Flow cytometry analysis of lipid peroxidation using the Cll-BODIPY probe. The histogram (left) shows a rightward shift in the LPS-treated group, indicating increased lipid peroxidation, which is reversed by FABP5 silencing. The violin plot (right) quantifies the Median Fluorescence Intensity (MFI). Data in (H) and (I) are presented as mean ±SEM (n>3 per group). Statistical significance was determined using a one-way ANOVA followed by Tukey’s post-hoc test. *p < 0.05, **p < 0.01.

Consistently, FABP5 expression strongly correlated with inflammatory mediators (*TNFRSF1B, CXCL2, ADAM17, JUNB*) and ferroptosis-related regulators (*LPCAT3, STEAP3, SAT1, SLC39A8*), while showing negative correlation with the survival-associated genes such as *PIK3R1* and *TSR2* (Fig. 2C). These relationships suggest that higher FABP5 levels accompany both inflammatory activation and compromised cell viability in MSA cerebellar white matter. Together, these data indicate that *FABP5* is functionally linked to pro-inflammatory signaling and ferroptosis in MSA. Based on these data and our co-expression findings, we propose that *FABP5* functions as a pathogenic bridge linking neuroinflammation and ferroptosis, thereby potentially contributing to oligodendrocyte loss in MSA (Fig. 2D).

### 3.3 FABP5 upregulation is specific to reactive astrocytes, supporting a non-cell-autonomous pathogenic mechanism

Our in-silico analysis strongly implicated FABP5 in a TNF/ferroptosis network (Fig. 2). However, MSA pathology is classically defined by GCI accumulation in oligodendrocytes. This presented a critical question: which cell type is the primary source of the upregulated FABP5? To address this, we first evaluated glial composition changes in MSA cerebellar white matter by performing cell-type deconvolution.Cell-type deconvolution of cerebellar white matter RNA-seq data (GSE199715) using scRNA-seq data (GSE67835) revealed notable shifts in glial composition:: astrocyte proportions were significantly increased, microglial fractions were reduced, while oligodendrocyte abundance showed no significant difference (Fig. 3A-C), partially supported by previous reports showing profound glial remodeling in MSA white matter [34, 35]. Notably, although microglial activation and proliferation are widely recognized pathological features of MSA [36], our scRNA-seq-based estimation involved a limited microglial sample size (n = 9), which may contribute to the observed discrepancy.

Single-cell RNA sequencing of mouse brains (SRA850958:SRS4386089) further clarified the cellular localization of FABP5. t-SNE visualization demonstrated that *Fabp5* was expressed in both oligodendrocyte and astrocyte clusters, suggesting a potential role in glial metabolic and inflammatory regulation (Fig. 3D). However, analysis of pathological tissue provided the definitive answer. Immunofluorescence staining confirmed a marked increase of FABP5 in the cerebellar white matter of PLP-α-syn mice, accompanied by expansion of GFAP-positive astrocytes (Fig. 3E-F), in agreement with previously reported glial activation in this model [37].

Most critically, FABP5 expression showed no significant changes in oligodendrocytes isolated from human MSA patients compared with controls (Fig. 3G-H). Multiplex immunofluorescence in the mouse model confirmed this human finding: FABP5 upregulation was specifically localized within GFAP-positive astrocytes (Fig. 3I-J). Together, these results demonstrate that FABP5 upregulation in MSA predominantly originates from astrocytes rather than oligodendrocytes, providing the first strong cellular-level evidence for a non-cell-autonomous pathogenic mechanism driven by reactive astrocytes.

### 3.4 FABP5 silencing mitigates inflammatory and ferroptotic responses in astrocytes

Our data thus far built a strong hypothesis: reactive astrocytes (Fig. 3) drive pathology by upregulating FABP5 (Fig. 1), which in turn orchestrates a TNF and ferroptosis gene signature (Fig. 2). To functionally validate this hypothesis, we examined whether astrocytic FABP5 participates in inflammatory and ferroptotic signaling. Primary astrocytes were isolated and purified, then divided into three groups: control cells, lipopolysaccharide (LPS)-treated cells to induce inflammatory environment [38], and LPS plus *Fabp5*-silenced (FABP5i) cells to assess the role of FABP5 in this context (Fig. 4A). FABP5 silencing was achieved using an shRNA plasmid (pRNAi-U6neo-FABP5), with Sanger sequencing and WB validation confirming efficient knockdown (Fig. S2A-D).

Astrocytes exposed to LPS displayed a distinct transcriptional shift that separated them from controls, while FABP5 knockdown produced a reciprocal shift toward baseline expression (Fig. 4B). LPS upregulated a network of pro-inflammatory and ferroptosis-related transcripts. Importantly, 1,471 genes were commonly modulated by LPS and reversed by FABP5 silencing, and pathway enrichment of these genes perfectly confirmed our in-silico hypothesis, highlighting TNF signaling and ferroptosis (Fig. 4C). Heatmap and volcano-plot views show robust induction of canonical inflammatory mediators (*Tnf*, *Tnfrsf1b*, *Adam17*) [39–41] and ferroptosis-linked genes (*Cybb*, *Tfrc*, *Gpx3*) [42–44] after LPS, and clear attenuation of these changes after FABP5 knockdown (Fig. 4D– E, 4G). Protein-network analysis indicates these LPS-responsive factors form a tightly connected module, consistent with a coordinated FABP5-dependent regulatory program (Fig. 4F).

We then validated these transcriptional findings at the protein and functional levels. Western blot analysis (Fig. 4H) provided direct protein-level confirmation. As suggested by the RNA-seq data, LPS treatment significantly induced the expression of the pro-inflammatory cytokine TNF-α and upregulated the ferroptosis-related protein GPX3 and ACSL5. Critically, FABP5i successfully blunted this induction, markedly reducing the protein levels of both TNF-α, GPX3, and ACSL5. Furthermore, to directly measure the functional endpoint of ferroptotic stress, we assessed lipid peroxidation using the C11-BODIPY probe. Flow cytometry analysis (Fig. 4I) revealed a significant increase in lipid ROS in LPS-treated astrocytes. This increase was completely prevented by *Fabp5* silencing. Together, these data provide robust functional confirmation that astrocytic *Fabp5* promotes an LPS-induced inflammatory state and the subsequent ferroptotic cascade.

### 3.5 FABP5 upregulation drives mitochondrial dysfunction in astrocytes

Finally, we sought to identify the ultimate downstream functional consequence of this *Fabp5*-driven inflammatory and ferroptotic state. Given that FABP5 promotes inflammatory and ferroptotic signaling, we next examined whether it contributes to mitochondrial impairment in astrocytes. Differential gene expression analysis revealed that LPS treatment upregulated *Tfrc* and *Cybb* while downregulating *Gpx4* and *Slc40a1*, indicating enhanced iron import, ROS generation, and weakened antioxidant defense (Fig. 5B). These transcriptional changes are consistent with a ferroptotic-like metabolic shift that increases oxidative burden[45]. Heatmap analysis of expression profiles confirmed this broad metabolic deregulation (Fig. 5C), LPS significantly elevated genes involved in lipid metabolism, ROS accumulation, and mitochondrial stress. Crucially, FABP5 silencing effectively reversed these transcriptional alterations, restoring the profiles toward control levels (Fig. 5C). The schematic model summarizes this mechanism, where LPS-induced FABP5 elevation facilitates PUFAs trafficking and amplifies mitochondrial oxidation imbalance through NOX2-derived ROS[46], collectively driving lipid peroxidation (Fig. 5A).

**Figure 5.**
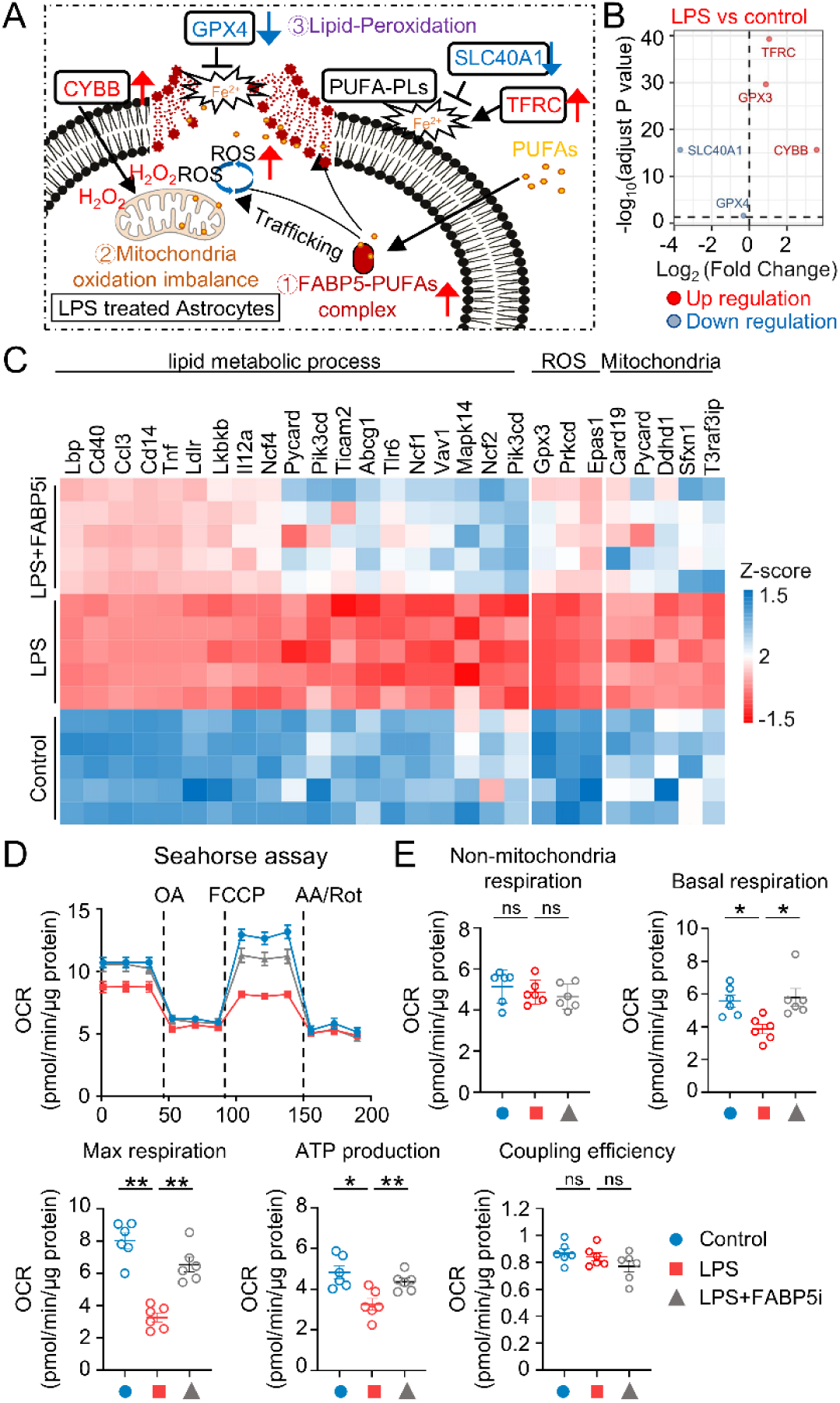
FABPS mediates LPS-induced mitochondrial dysfunction in astrocytes. (A) Schematic of the proposed mechanism. Lipopolysaccharide (LPS) stimulation upregulates FABP5, which facilitates the trafficking of polyunsaturated fatty acids (PUFAs). This process, combined with upregulated ROS production from CYBB (NOX2) and mitochondrial oxidation imbalance, drives excessive lipid peroxidation. The ferroptotic cascade is amplified by the simultaneous downregulation of key defense proteins GPX4 and the iron exporter SLC40Al, alongside the upregulation of the iron importer TFRC. (B) Volcano plot of differentially expressed genes (DEGs) from RNA-sequencing analysis comparing LPS-treated primary astrocytes with vehicle-treated controls. Key ferroptosis regulators (Tfrc, Cybb, Gpx4, Slc40al) are highlighted. Red dots indicate significantly upregulated genes; blue dots indicate significantly downregulated genes. (C) Heatmap displaying the Z-scores of DEGs related to lipid metabolic processes, ROS, and mitochondrial pathways in astrocytes under Control, LPS, and LPS + FABP5i conditions. (D) Representative trace of oxygen consumption rate (OCR) measured by a Seahorse analyzer in astrocytes treated as indicated. Sequential injections of oligomycin (OA), FCCP, and antimycin A/roteneone (AA/Rot) are marked by dashed lines. (E) Quantification of non-mitochondrial respiration, basal respiration, maximal respiration, ATP production, and coupling efficiency derived from the Seahorse assay in (D). Data in (D) and (E) are presented as mean ±pm$ SEM (n=5-6 per group). Statistical significance was determined using a one­ way ANOVA followed by Tukey’s post-hoc test. *p < 0.05, **p < 0.01; ns, not significant.

To validate the functional impact on cellular energy, we performed Seahorse XF analysis (Fig. 5D-E). LPS exposure resulted in a profound reduction in oxygen consumption rate (OCR), confirming impaired mitochondrial respiration. Conversely, FABP5i significantly restored key parameters of mitochondrial function, including basal respiration, maximal respiration, ATP production, and coupling efficiency, while reducing non-mitochondrial OCR (Fig. 5E). Together, these findings confirm that FABP5 acts as a critical link between inflammation-induced oxidative stress and mitochondrial dysfunction in astrocytes. This final step provides the complete mechanistic link: from FABP5 upregulation to TNF/ferroptosis signaling and culminating in the functional mitochondrial collapse that defines the pathology.

## 4. Discussion

Glial cell inclusions of the oligodendroglia and cerebellar white matter dysfunction is a central pathological hallmark of (MSA, yet the underlying molecular pathogenesis remain poorly understood [47, 48]. The present study provides a comprehensive and novel mechanistic framework for white matter degeneration in MSA. We identify astrocytic FABP5 as a critical and previously unrecognized pathogenic mediator. Our data, for the first time, mechanistically link three key processes: dysregulated lipid metabolism (mediated by FABP5), TNF-driven neuroinflammation, and ferroptotic oxidative stress. We demonstrate that this pathological cascade ultimately converges on a catastrophic failure of mitochondrial function (Fig. 6).

**Figure 6.**
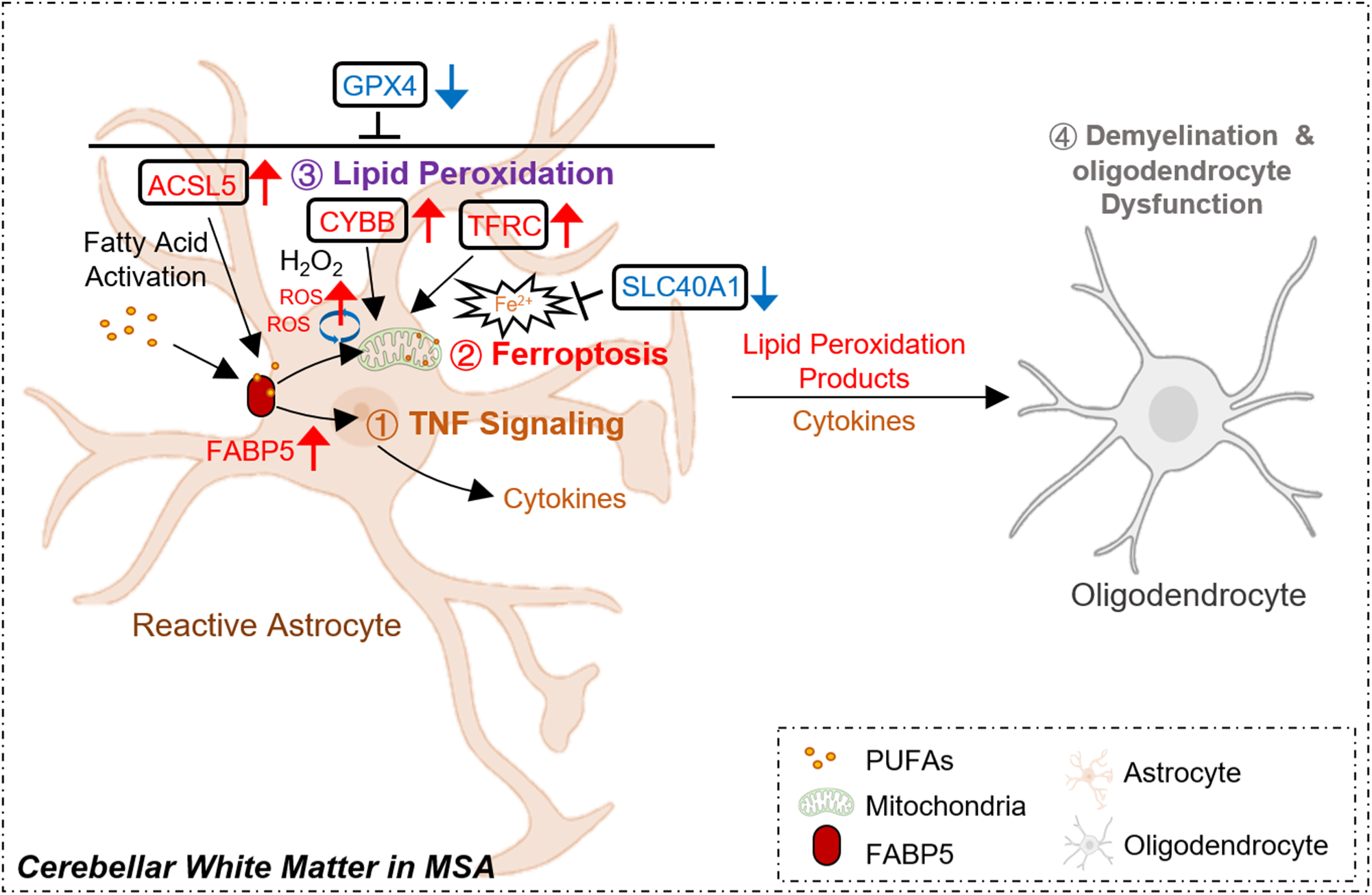
Schematic of the proposed non-cell-autonomous mechanism driving oligodendrocyte dysfunction in MSA. In the cerebellar white matter of MSA, reactive astrocytes become a primary driver of pathology. (1) Upregulated astrocytic FABP5 acts as a central mediator, facilitating the trafficking of PUFAs and orchestrating two pathogenic pathways: (1) the activation ofTNF signaling, leading to the release of pro-inflammatory TNF-aor other cytokines; and (2) the initiation of ferroptosis. (3) This ferroptotic state is characterized by the upregulation of ACSL5 (for fatty acid activation), CYBB (generating ROS), and TFRC (increasing intracellular iron Fe^2^+), coupled with the downregulation of key ferroptosis defenses GPX4 and the iron exporter SLC40Al. This cascade results in excessive lipid peroxidation. (4) Finally, the combination of inflammatory mediators and oxidative stress signals exported by the reactive astrocyte leads to the dysfunction of neighboring oligodendrocytes, driving white matter pathology.

The most transformative contribution of this work is the reframing of MSA pathology from a classic cell-autonomous model[49] to one driven by a non-cell-autonomous mechanism. MSA is pathologically defined by the accumulation of GCIs within oligodendrocytes[50], a finding that has long implied a “self-destructive” process. Our data fundamentally challenge this dogma. We demonstrate that the key pathogenic driver, FABP5, is not upregulated in the “victim”, the oligodendrocyte (Fig. 3G, 3H), but is instead dramatically overexpressed in the “attacker”, the neighboring reactive astrocyte (Fig. 3E, 3I, 3J). This critical spatial evidence recasts astrocytes not as passive bystanders to α-synuclein pathology, but as toxic effectors that actively drive oligodendrocyte demise.

Transcriptomic profiling of MSA cerebellar white matter revealed a robust activation of inflammatory and ferroptotic pathways, most prominently involving TNF and VEGF signaling. Such glial activation is consistent with the chronic neuroinflammatory milieu reported across synucleinopathies[51]. In MSA, α-synuclein pathology disrupts glial homeostasis, leading to cytokine release and reactive gliosis that contribute to demyelination[12]. Crucially, the concomitant upregulation of ferroptosis-related genes, including *SLC39A8*, *STEAP3*, and *SAT1*, suggests a redox-dependent component of glial injury[52–54]. These findings indicate that inflammation and ferroptosis do not act in isolation but instead jointly constitute a pathogenic axis in MSA.

Our co-expression network analysis identified FABP5 as the central metabolic hub linking the enriched inflammatory and ferroptotic modules. FABP5 is an established cytosolic lipid chaperone regulating long-chain fatty acid (LCFA) transport and nuclear signaling via pathways like NF-κB and PPAR[55–57]. By demonstrating that FABP5 expression is highly correlated with both TNF-related and ferroptosis genes, our results extend prior findings in PD and ischemic models, which showed that FABP5 amplifies inflammatory gene expression and oxidative stress[25, 58]. This finding provides the first evidence that FABP5 serves as a pivotal node for linking lipid peroxidation with inflammatory signaling in MSA glial cells, thereby initiating the chronic redox imbalance in MSA white matter. Furthermore, the negative correlation with survival-associated genes like *PIK3R1* (Fig. 1E, 2C) suggests a potential’dual-hit’ mechanism, whereby FABP5 not only promotes toxic-gain-of-function (inflammation, ferroptosis) but also contributes to a loss-of-protection (impaired PI3K/AKT signaling).

Previous studies on FABP5 further support its involvement in α-syn oligomerization[59] and glial stress[29]. The dual inhibitory effect of *Fabp5* silencing on LPS-induced pro-inflammatory cytokine release and MSA-associated ferroptosis rigorously validates its role as a key orchestrator of meta-inflammation. We hypothesize that the dysregulation of FABP5 disrupts the cellular handling of PUFAs, leading to their toxic accumulation and increased susceptibility to oxidative damage, a hallmark of ferroptosis. By demonstrating that *Fabp5* silencing restores ferroptosis genes like *Cybb*, *Tfrc* and *Gpx3* function while inhibiting cytokine release, we establish FABP5 as the metabolic driver linking lipid peroxidation and inflammation. This targeted disruption represents a highly promising strategy to break the self-propagating cycle of chronic neuroinflammation that accelerates MSA progression.

Functional validation shows that *Fabp5* silencing profoundly protects mitochondrial integrity and promotes cell survival in astrocytes. This suggests FABP5 has a direct role in the cell death mechanism. This protective effect aligns with molecular studies showing that FABP5 accelerates cell death by mediating mitochondrial outer membrane permeabilization[24]. Mechanistically, FABP5’s interaction with VDAC-1 and BAX is known to facilitate the formation of mitochondrial macropores[24]. Therefore, our findings position astrocytic FABP5 as the convergence point where the pathological inflammatory and ferroptotic signals merge. This merging triggers the lethal mitochondrial breakdown that may subsequently impact vulnerable oligodendrocytes, establishing FABP5 as an optimal therapeutic target to preserve MSA white matter.

Although this study reveals a pivotal role of astrocytic FABP5 in mediating inflammatory, ferroptotic, and mitochondrial injury in MSA, several limitations warrant consideration. First, our findings are based primarily on bulk and single-cell transcriptomic analyses and *in vitro* validation using LPS-induced astrocyte models. While these approaches effectively capture the molecular landscape, they cannot fully reproduce the complex *in vivo* microenvironment of MSA, where multiple glial and neuronal populations interact dynamically. Second, although FABP5 upregulation was confirmed in both human and mouse tissues, its causal relationship with α-syn aggregation or oligodendrocyte loss remains to be mechanistically dissected. Third, our deconvolution analysis (Fig. 3C) showed a surprising decrease in microglia, which, as noted in the results, is likely a technical artifact of the small microglial sample size (n=9) in the reference dataset (GSE67835) and requires further validation. Finally, a definitive *in vivo* rescue experiment—such as astrocyte-specific *Fabp5* knockdown in the PLP-α-syn mouse model to assess oligodendrocyte preservation and motor outcomes—is required to fully confirm the therapeutic potential of this target. Given FABP5’s dual role in lipid metabolism and inflammation, future work should integrate metabolomic and lipidomic profiling to provide deeper insight into its contribution to ferroptotic vulnerability and energy dysregulation in MSA.

These findings also open several new avenues for translational research. As a’druggable’ intracellular lipid chaperone, FABP5 is an exceptionally attractive target. Small molecule inhibitors, some of which have shown efficacy in other demyelinating disease models, represent a direct translational path[60–62]. It also remains to be determined if FABP5 itself, or its downstream lipid-peroxidation products, can be measured in CSF as a biomarker to track disease progression or target engagement. Finally, this work raises a broader question: Is this astrocytic FABP5-driven’Astrocyte-Ferroptosis-Mitochondrial axis’ a common pathogenic pathway in other synucleinopathies or primary white matter disorders?

In summary, our study identifies FABP5 as a central astrocytic mediator linking lipid metabolism, inflammation, ferroptosis, and mitochondrial dysfunction in multiple system atrophy. By integrating transcriptomic profiling, co-expression analysis, and functional validation, we demonstrate that FABP5 upregulation drives proinflammatory signaling and ferroptotic stress, ultimately compromising astrocytic metabolic integrity and glial homeostasis. Silencing FABP5 not only alleviates inflammatory responses but also restores redox and mitochondrial balance, highlighting its dual role as a metabolic and immune regulator. These findings advance our understanding of glial pathology in MSA and propose FABP5 as a promising therapeutic target within the oxidative-metabolic axis of neurodegeneration.

## CRediT authorship contribution statement

C.W., and J.L.: original draft writing, coding, visualization, validation, data curation. Y.C.: coding and data analysis. N.T.: *in vivo* study., Q.C., B.L., F.D., and W.Z.: data curation. D.F.: supervision, and the PLP-α-syn transgenic mouse. I.K., K.F., and A.C.: funding acquisition. A.C.: writing-review & editing, supervision, funding acquisition, conceptualization. All authors critically reviewed and approved the final version of the manuscript. A.C. verified the data and had the final responsibility for the decision to submit for publication.

## Acknowledgments

This work was supported by the Start-up Fund from Sun Yat-sen University (Grant No. 59000-12256011) and the General Program of Shenzhen Natural Science Foundation (Grant No. JCYJ20250604174755073). This research was also supported in part by the Japan Agency for Medical Research and Development (AMED) (Grant Nos. 24ym0126095h0003, 25ym0126189h0001); the Japan Science and Technology Agency (JST) (Grant Nos. MASP202432, JPMJSF2512); the Japan Society for the Promotion of Science (JSPS) KAKENHI (Grant Nos. 22K06644, 25K10004); and The Takeda Science Foundation. The authors wish to thank Gemini 2.5 Pro for assistance with English language polishing of this manuscript.

## Data availability

The bulk RNA-sequencing data generated in this study for the primary astrocyte experiments (Fig. 4, 5) have been deposited in the NCBI Gene Expression Omnibus (GEO) and are available under the accession number (In coming). The publicly available datasets analyzed during the current study are available in the GEO repository under accession numbers GSE199715 (human MSA cerebellar white matter RNA-seq), GSE199724 (human laser-captured oligodendrocytes), and GSE67835 (reference scRNA-seq for deconvolution)[30]. The mouse brain single-cell RNA-seq dataset used for t-SNE visualization (Fig. 3) is available in the Sequence Read Archive (SRA) under accession SRA850958:SRS4386089[63]. All other data supporting the findings of this study are available within the article and its supplementary materials or from the corresponding author upon reasonable request.

## Supplementary materials

**Figure S1.**
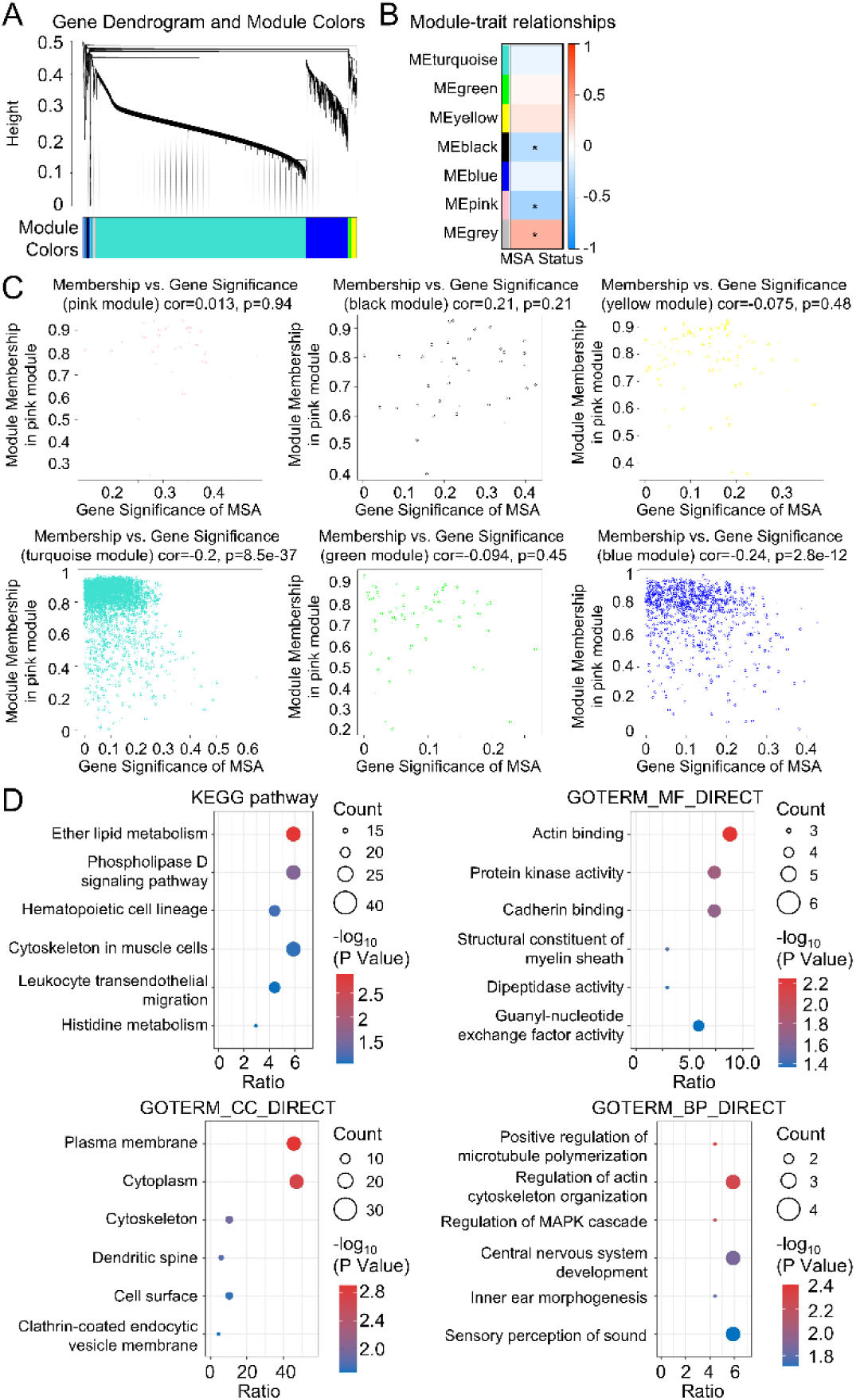
Weighted Gene Co-expression Network Analysis (WGCNA) of gene expression data. (A) Gene dendrogram and module colors. Genes were clustered using hierarchical clustering based on topological overlap, and co-expression modules were identified using the dynamic tree cut method. Each color represents a distinct module. (B) Module - trait relationship heatmap. The heatmap displays the Pearson correlation between each module eigengene (ME, representing the overall expression profile of a module) and the MSA disease status. Red indicates a positive correlation, and blue indicates a negative correlation. The numbers in each cell represent the correlation coefficient (top) and the p-value (bottom, in parentheses). The pink and black modules show the most significant negative correlation with MSA status. (C) Scatterplots of module membership (MM) versus gene significance (GS) for each identified module. Each plot visualizes the correlation between how “central” a gene is to its module (MM, x-axis) and how strongly that gene is associated with MSA Krankheit (GS, y-axis). A significant correlation in this plot (e.g., as seen in the turquoise and blue modules) indicates a strong internal structure where hub genes are also highly significant for the disease, validating the biological relevance of the module’s construction.

**Figure S2.**
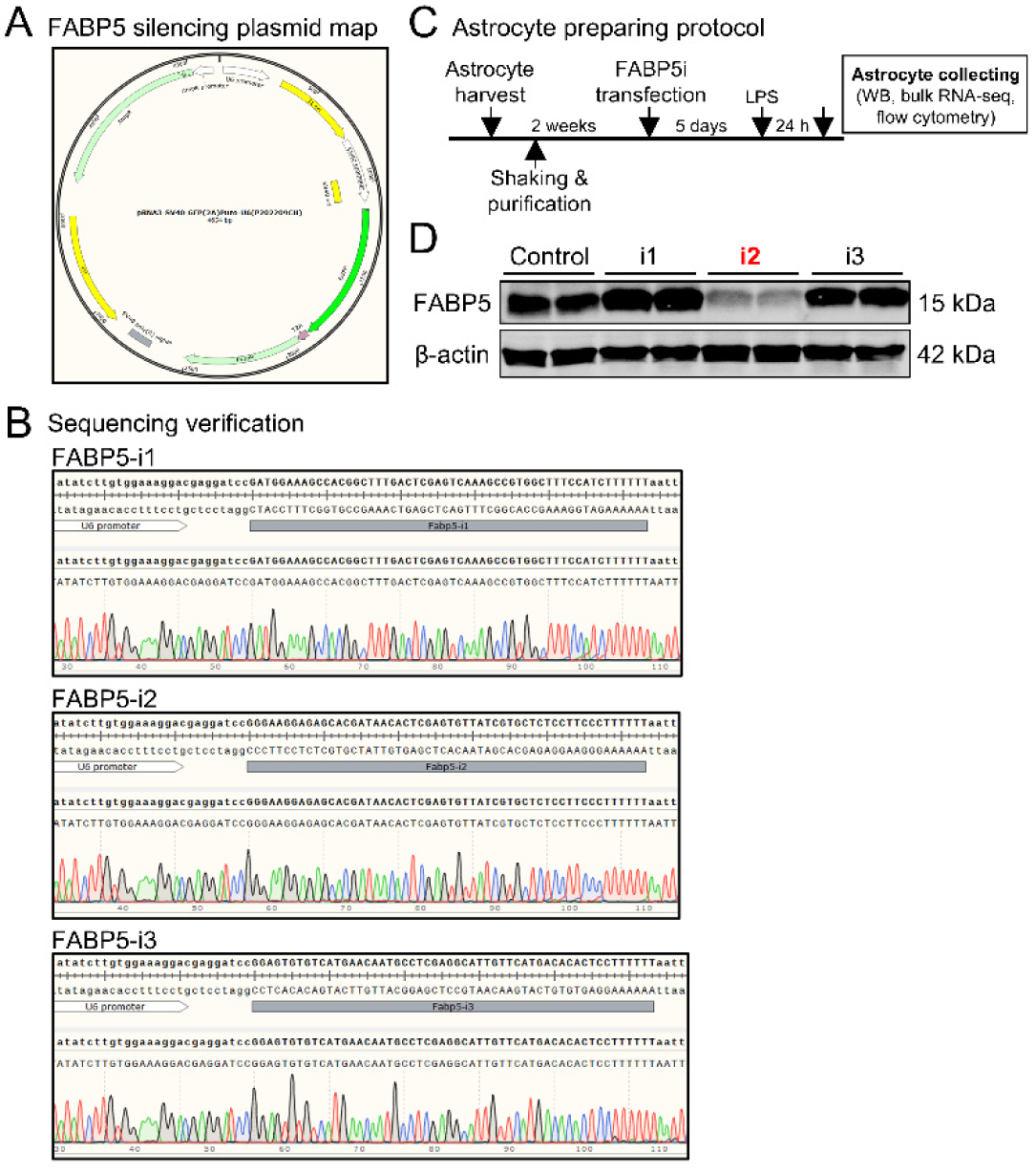
Generation and validation of FABPS silencing plasmids. (A) Plasmid’ map of the pRNAI-U6neo-F ABP5(shRNA) construct used for silencing mouse FABP5 gene. (B) Sanger sequencing verification of the three different FABP5 shRNA’ constructs (FABP5-il, FABP5-i2, and FABP5-i3). The chromatograms confirm the correct insertion of the shRNA sequences downstream of the U6 promoter.(C) Schematic diagram illustrating the experimental protocol. Primary astrocytes were harvested, purified for 2 weeks, and then transfected with the FABP5-silencing (FABP5i) plasmids for 5 days. Cells were subsequently treated with LPS for 24 hours before being collected for Western blot (WB), bulk RNA-seq, and flow cytometry analyses.(D) Western blot analysis showing the knockdown efficiency of the three FABP5 shRNA constructs. FABP5 protein levels (∼15 kDa) are shown for Control, FABP5-il, FABP5-i2, and FABP5-i3 transfected cells. *f3*-actin (∼50 kDa) was used: as a loading control. The construct i2 (highlighted in red) demonstrates the most effective knockdown ofFABP5.

---

**Table 1.** Key resources.

| REAGENT & RESOURCE | SOURCE | IDENTIFIER | RRID |
| --- | --- | --- | --- |
| Antibodies |  |  |  |
| Mouse Anti-Beta Actin | Huabio | Cat. No. EM21002 | AB_2819164 |
| Mouse Anti-GFAP | Proteintech | Cat. No. 60190-1-Ig | AB_10838694 |
| Mouse Anti-GPX3 | Proteintech | Cat. No. 3947-1-AP | AB_3085426 |
| Rabbit Anti-ACSL5 | Proteintech | Cat. No. 15708-1-AP | AB_2257881 |
| Rabbit Anti-FABP5 | Proteintech | Cat. No. 12348-1-AP | AB_2100341 |
| Rabbit Anti-NOX2 | Proteintech | Cat. No. 19013-1-AP | AB_2833044 |
| Rabbit Anti- TNF Alpha | Proteintech | Cat. No. 17590-1-AP | AB_2271853 |
| Goat anti-mouse IgG (H+L), HRP conjugate | Proteintech | Cat. No. SA00001-1 | AB_2722565 |
| Goat anti-rabbit IgG (H+L), HRP conjugate | Proteintech | Cat. No. SA00001-2 | AB_2722564 |
| Donkey Anti-Mouse IgG H&L (Alexa Fluor® 568) | Abcam | Cat. No. ab175472 | AB_2636996 |
| Chemicals, peptides, and recombinant proteins |  |  |  |
| Gibco™ BASIC DMEM, High Glucose, Pyruvate | Thermo Fisher Scientific | Cat. No. C11995500BT |  |
| Penicillin/streptomycin | Thermo Fisher Scientific | Cat. No. 15140 |  |
| Fetal bovine serum | Enkilife | Cat. No. CMS0003 |  |
| Trypsin-EDTA (0.25%), phenol red | Thermo Fisher Scientific | Cat. No. 25200072 |  |
| 0.5M EDTA, pH8.0<br>(Sterile, Cell Culture Grade) | Beyotime | Cat. No. C0196 |  |
| PBS, 1× (pH7.4) | Servicebio | Cat. No. G4202 |  |
| Opti-MEM™ I Reduced Serum Medium | Thermo Fisher Scientific | Cat. No. 31985070 |  |
| Lipopolysaccharide (LPS) | Sigma-Aldrich | Cat. No. L2630 |  |
| RIPA lysis buffer | Beyotime | Cat. No. P0013D |  |
| Protease Inhibitor Cocktail (EDTA-Free, 100× in DMSO) | APExBIO | Cat. No. K1007 |  |
| Phosphatase Inhibitor Cocktail 1 (100× in DMSO) | APExBIO | Cat. No. K1012 |  |
| 5× SDS loading buffer | Epizyme | Cat. No. LT101 |  |
| 10× Tris-glycine SDS-PAGE running buffer | Servicebio | Cat. No. G2027 |  |
| 10× transfer buffer | Servicebio | Cat. No. G2154 |  |
| Tween-20 | BioFroxx | Cat. No. 1247ML100 |  |
| 10× TBS | Servicebio | Cat. No. G0015 |  |
| Skim milk powder | BioFroxx | Cat. No. 1172GR500 |  |
| Multicolor Prestained Protein Ladder | Epizyme | Cat. No. WJ103 |  |
| Triton X-100 | BioFroxx | Cat. No. 1139ML100 |  |
| BSA albumin fraction V | BioFroxx | Cat. No. 4240GR100 |  |
| Fluoroshield mounting medium | Beyotime | Cat. No. P0128M |  |
| DAPI | Beyotime | Cat. No. C1006 |  |
| Critical commercial assays |  |  |  |
| E.Z.N.A.® Endo-Free Plasmid DNA Maxi Kit | Omega Bio-tek | Cat. No. D6926-01 |  |
| Efficient chemiluminescence kit | Absin | Cat. No. abs9434 |  |
| 12.5% PAGE gel rapid preparation kit | Epizyme | Cat. No. PG113 |  |
| 15% PAGE gel rapid preparation kit | Epizyme | Cat. No. PG114 |  |
| Lipofectamine™ LTX Reagent with PLUS™ Reagent | Thermo Fisher Scientific | Cat. No. 15338100 |  |
| Lipid peroxidation sensor (BODIPY 581/591 C11) | Beyotime | S0043S |  |
| E.Z.N.A.® Endo-Free Plasmid DNA Maxi Kit | Omega Bio-tek | Cat. No. D6926-01 |  |
| Efficient chemiluminescence kit | Absin | Cat. No. abs9434 |  |
| Experimental models: Organisms/strains |  |  |  |
| C57BL/6J mice | Jackson labs | Strain #: 000664<br>RRID:IMSR_JAX:000664 |  |
| Plp-αsyn mice | Jackson labs | | |
| Software and algorithms |  |  |  |
| GraphPad Prism | GraphPad Prism®, La Jolla, CA |  |  |
| R (Versions 3.6.2–4.1.1) | The R Foundation | https://www.r-project.org/foundation/;<br>RRID:SCR_001905R |  |
| Adobe Illustrator | Adobe, San Jose, CA |  |  |
| Photoshop | Adobe, San Jose, CA |  |  |
| Microsoft Excel | Microsoft Corporation<br>, Albuquerque, NM |  |  |

Code:

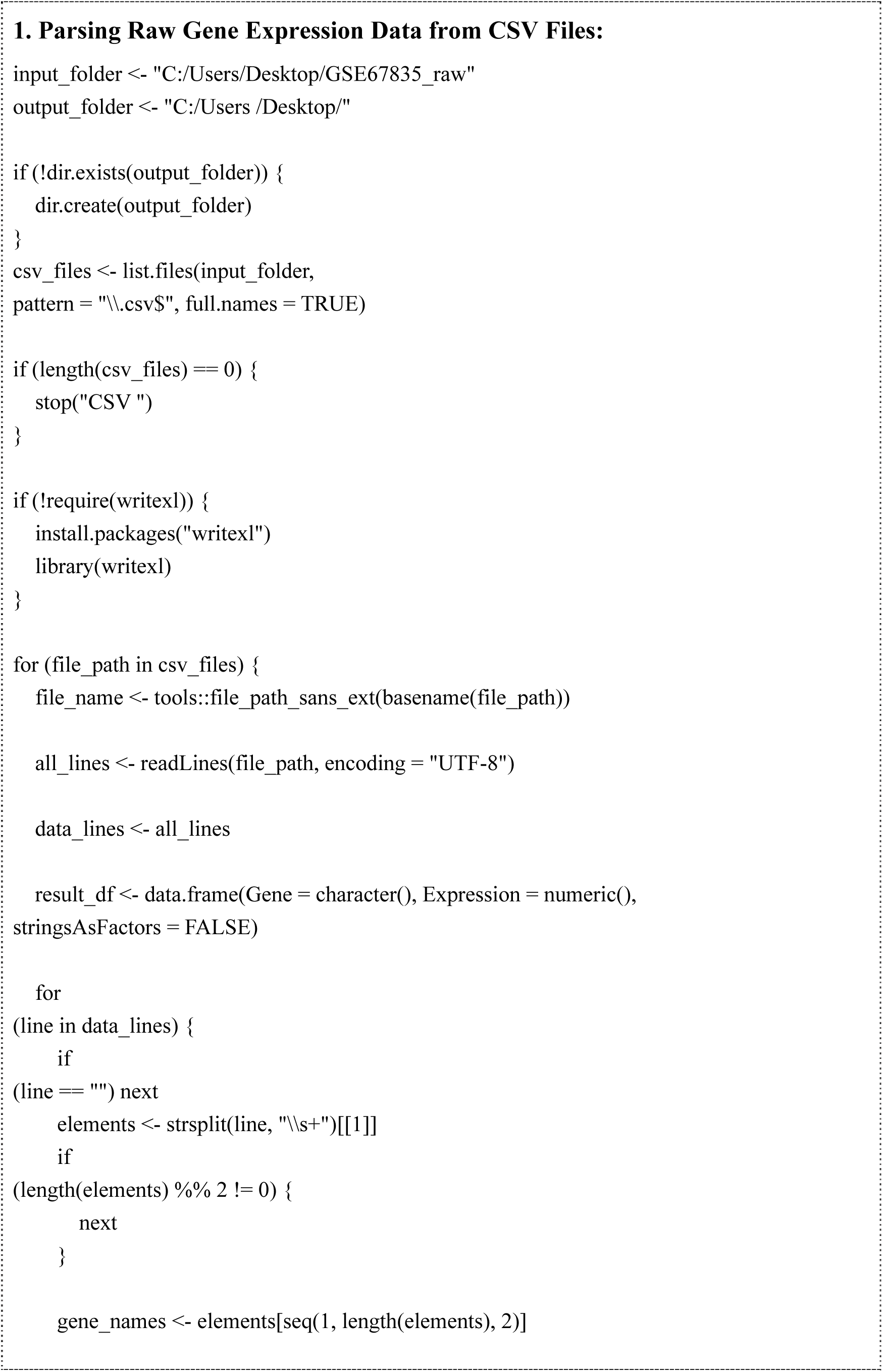
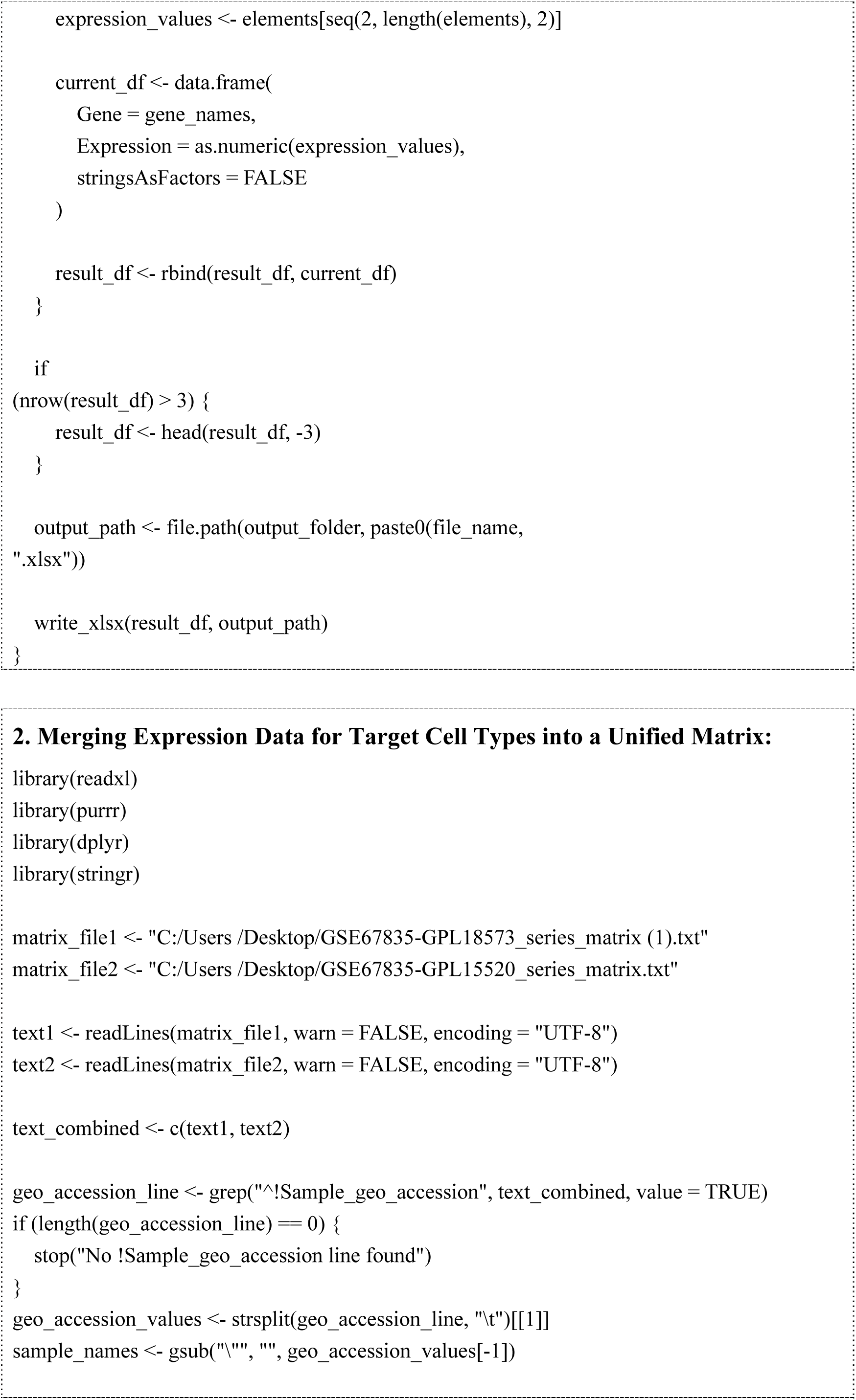
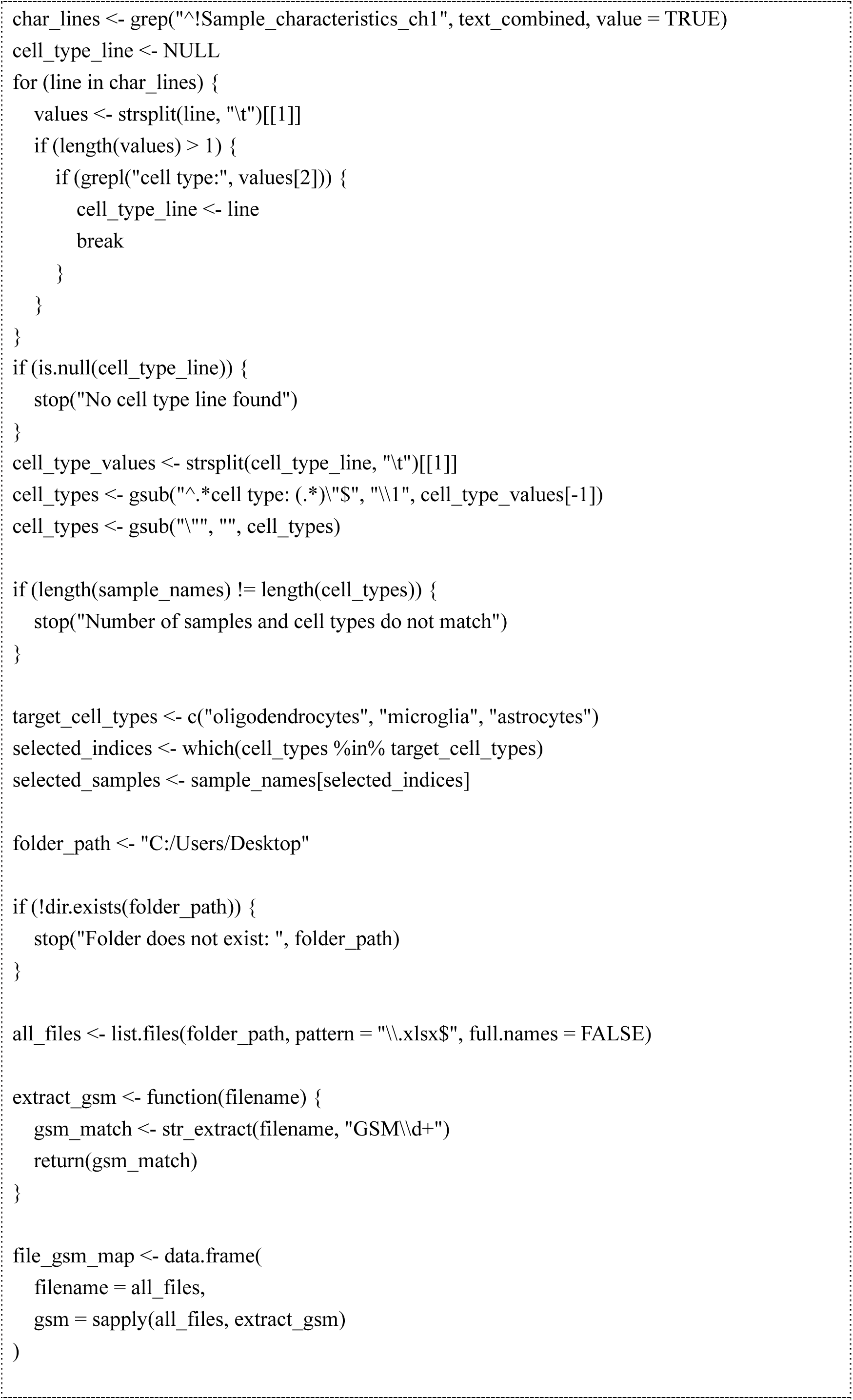
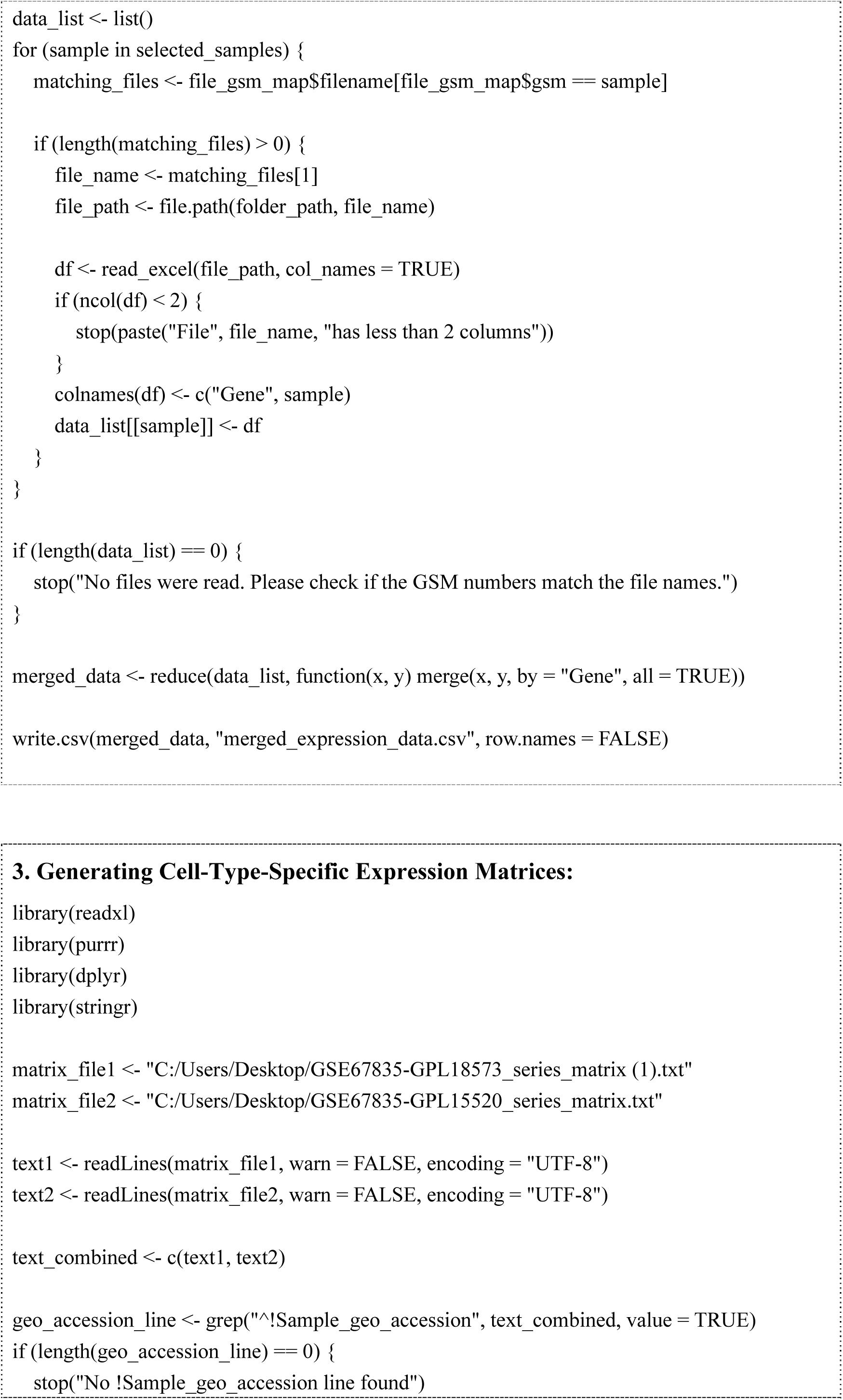
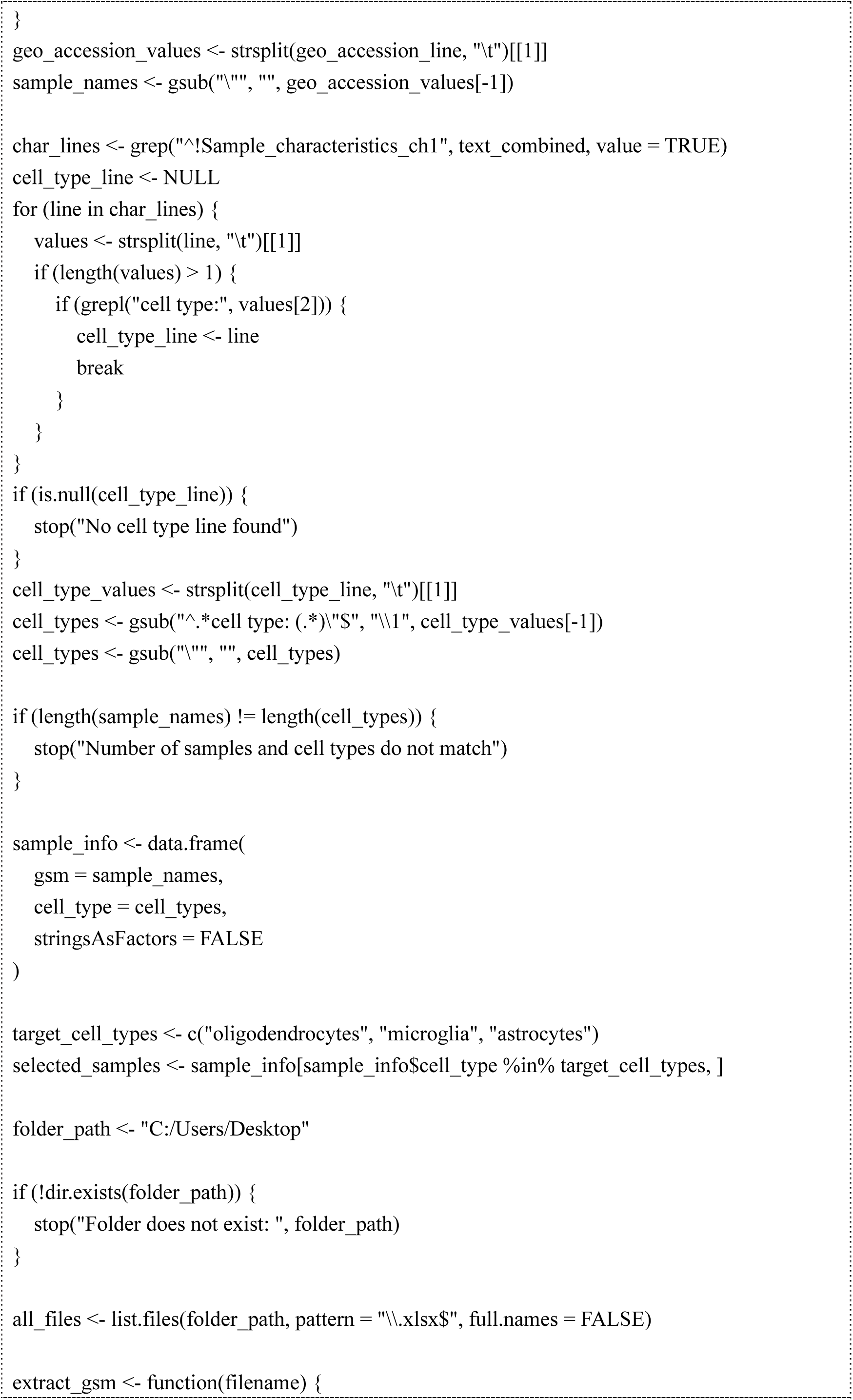
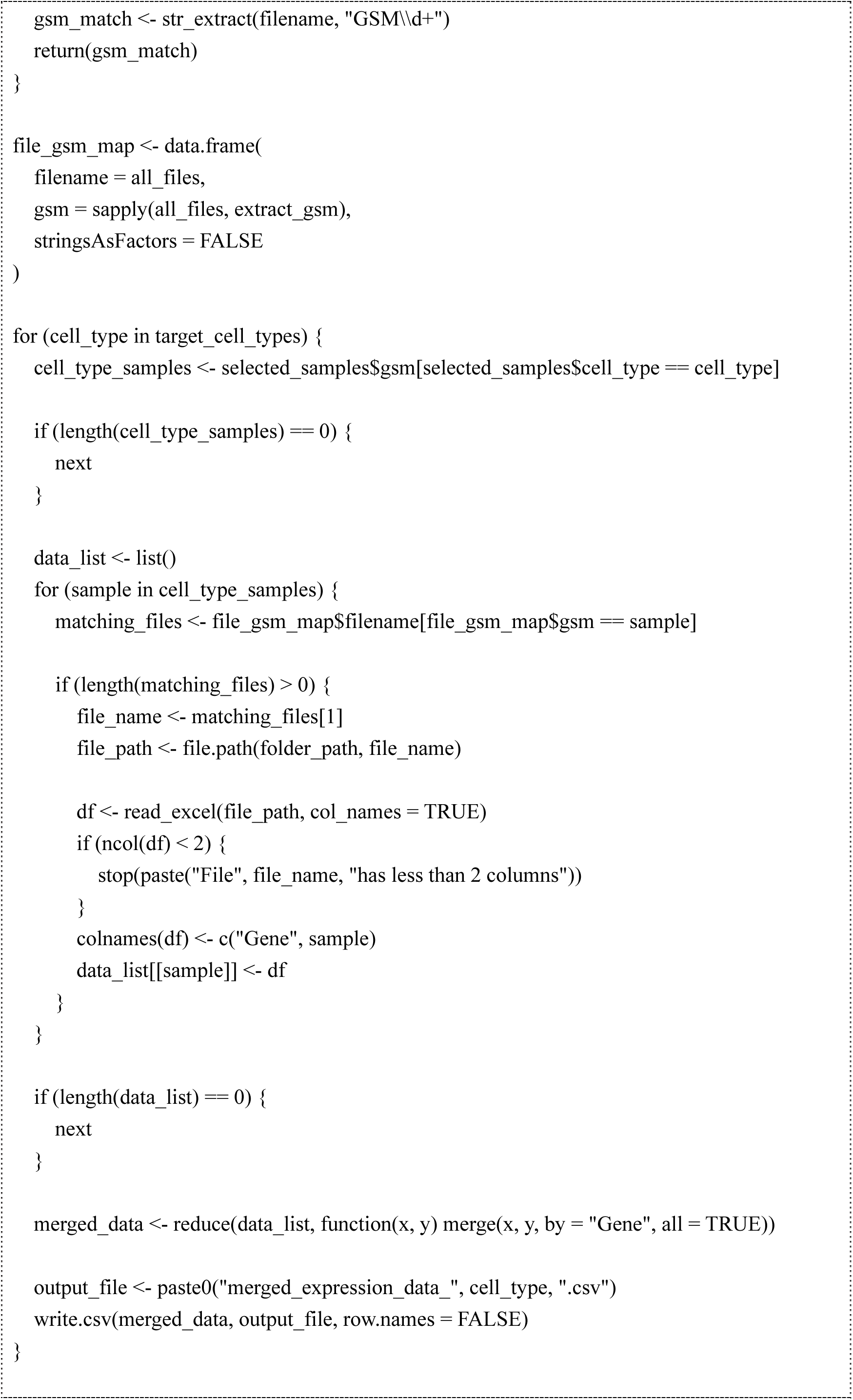
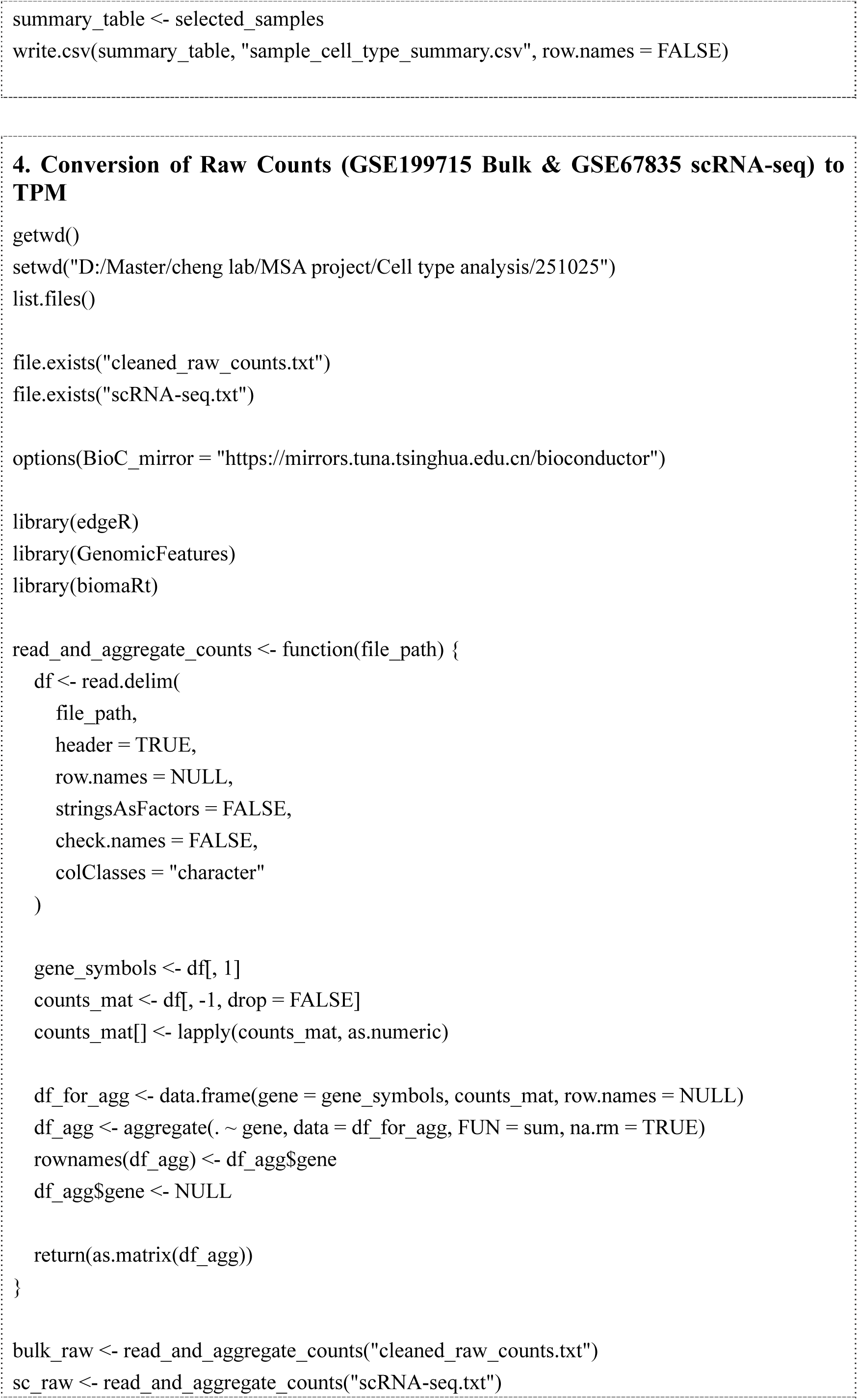
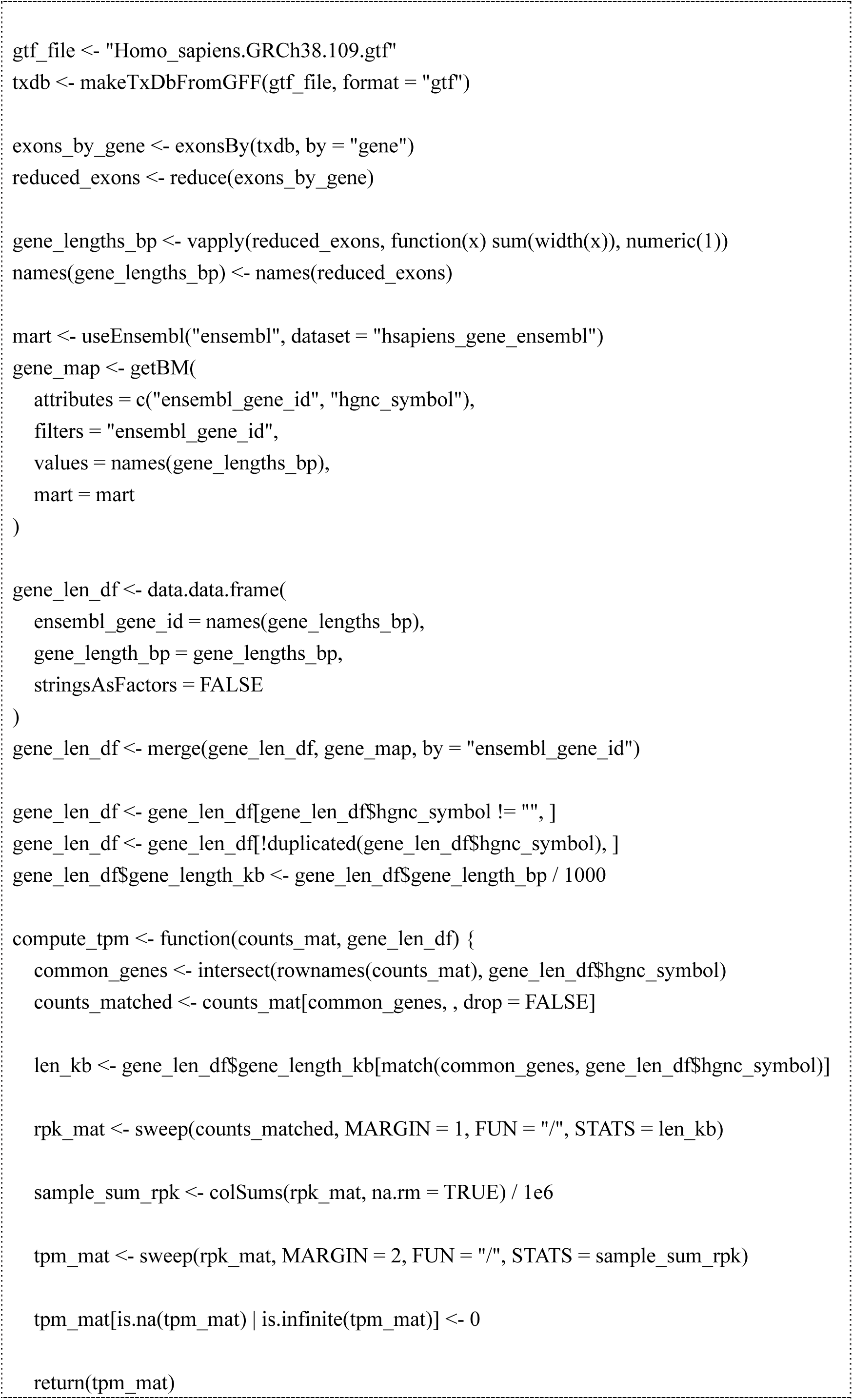
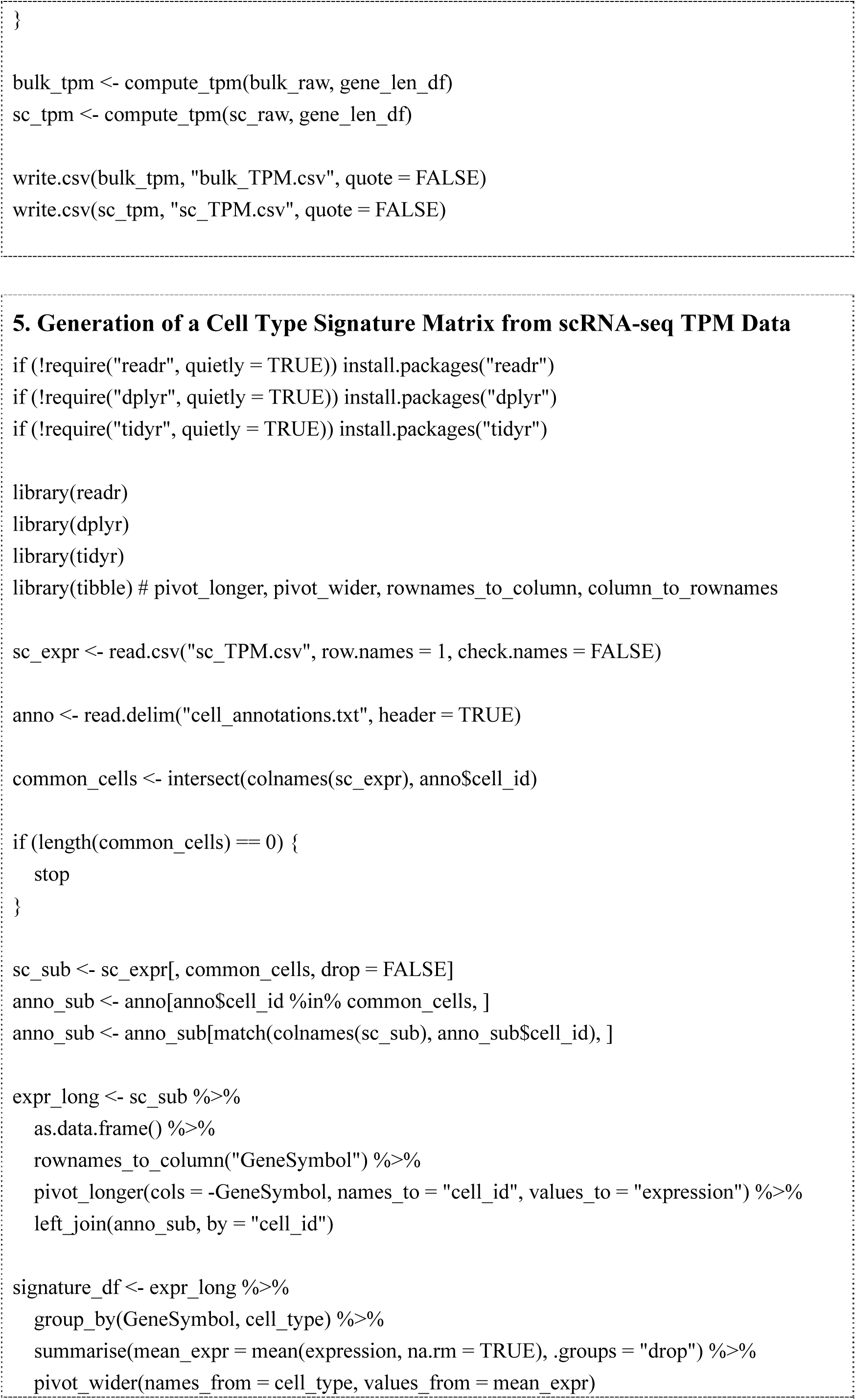
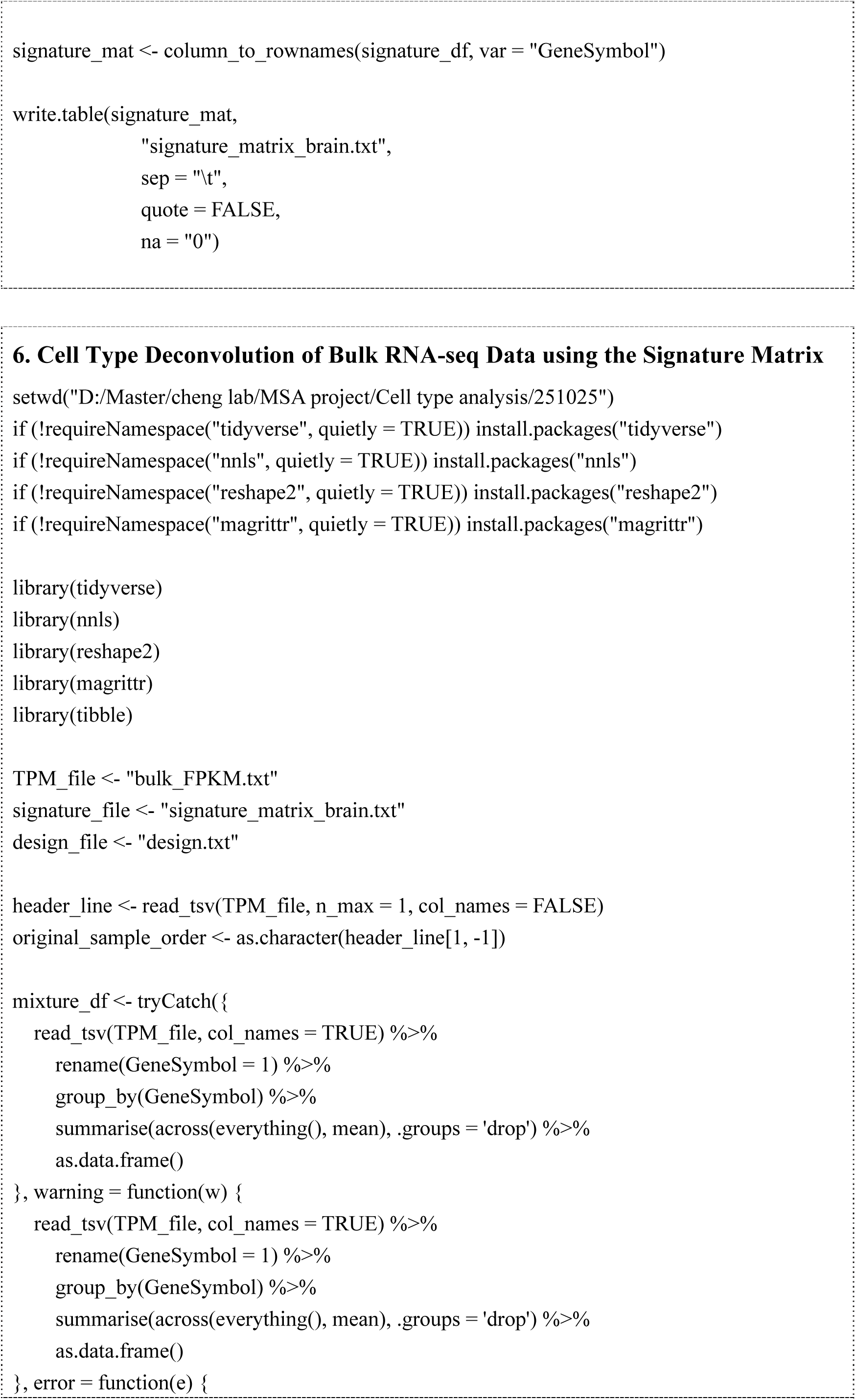
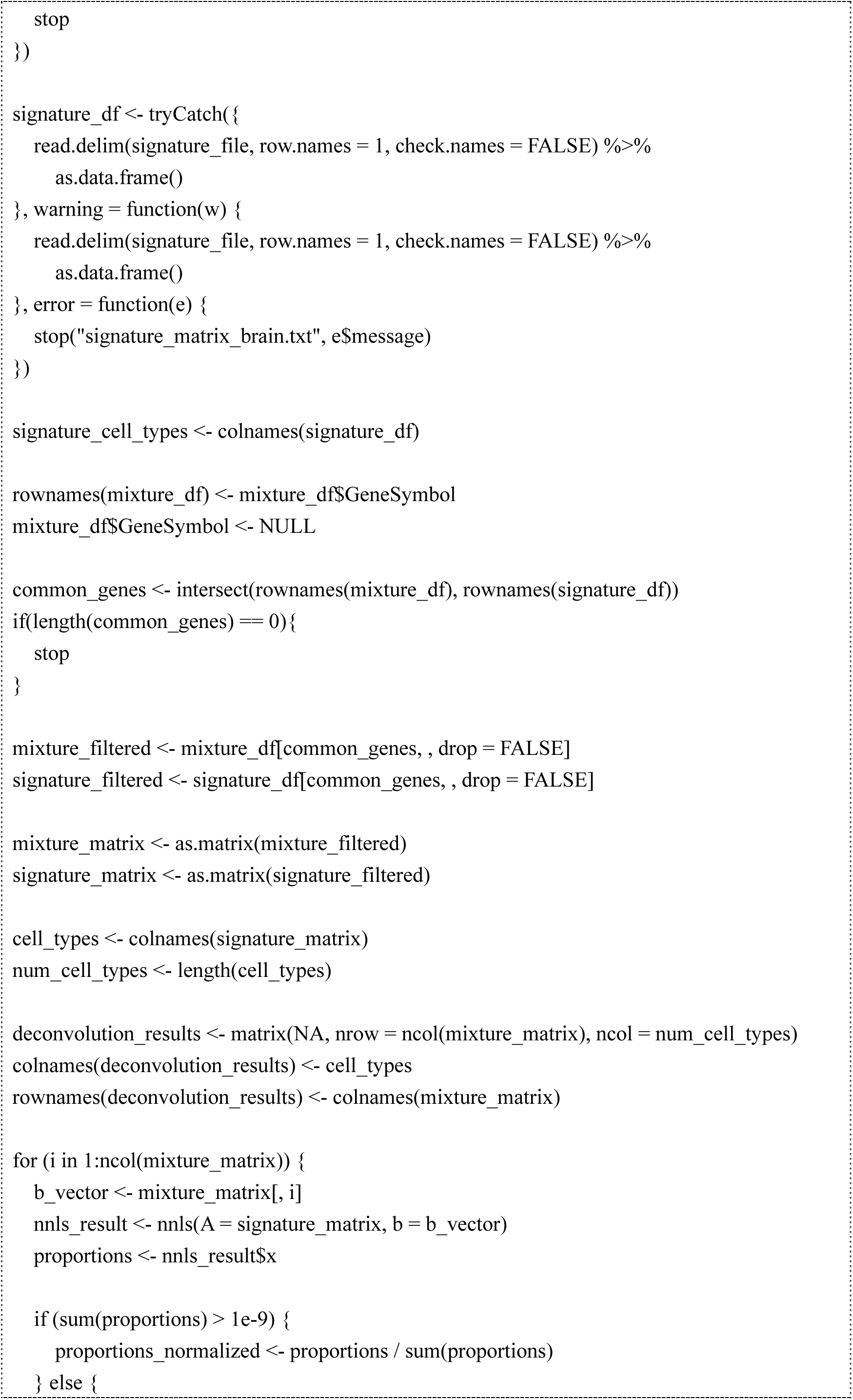
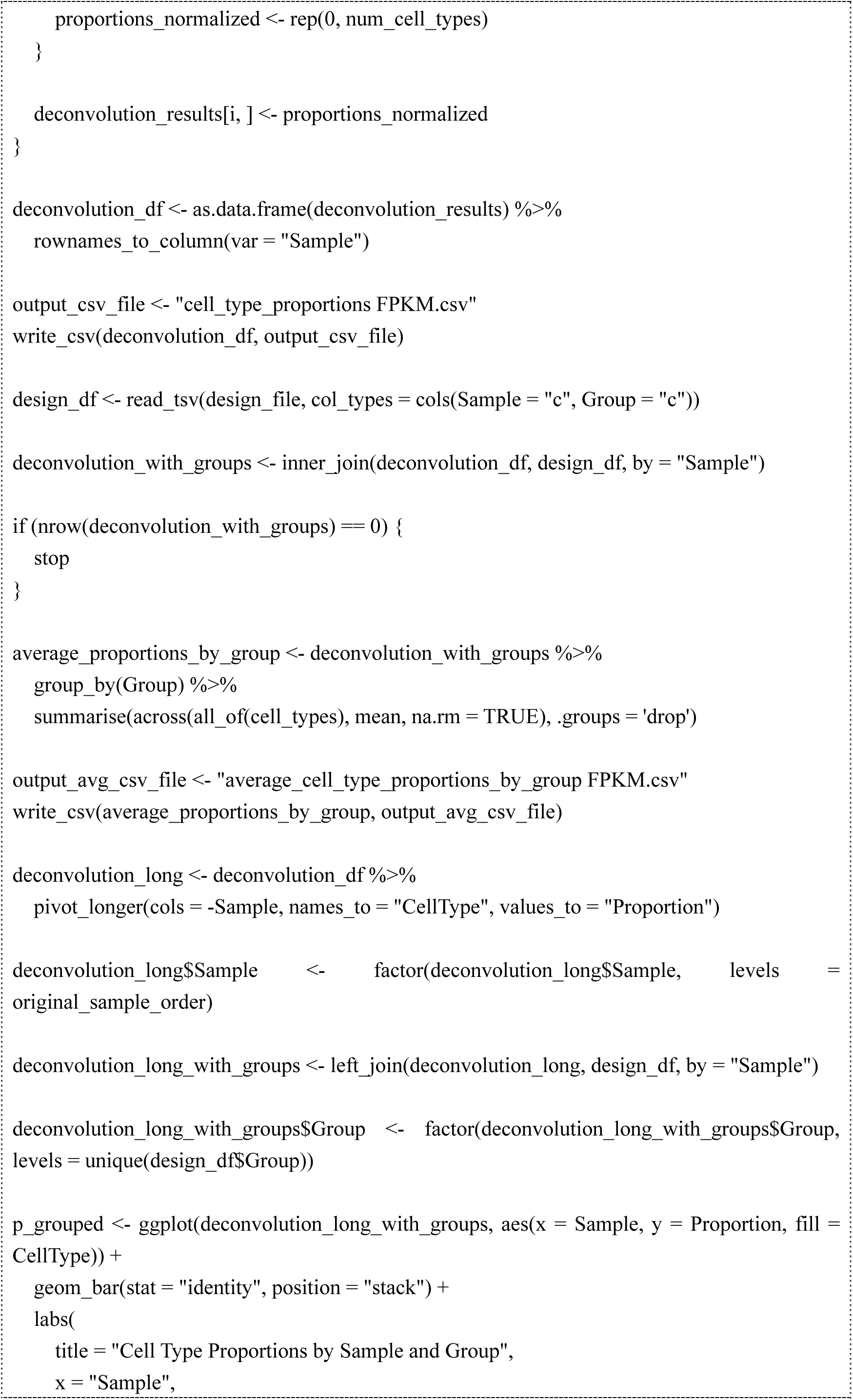
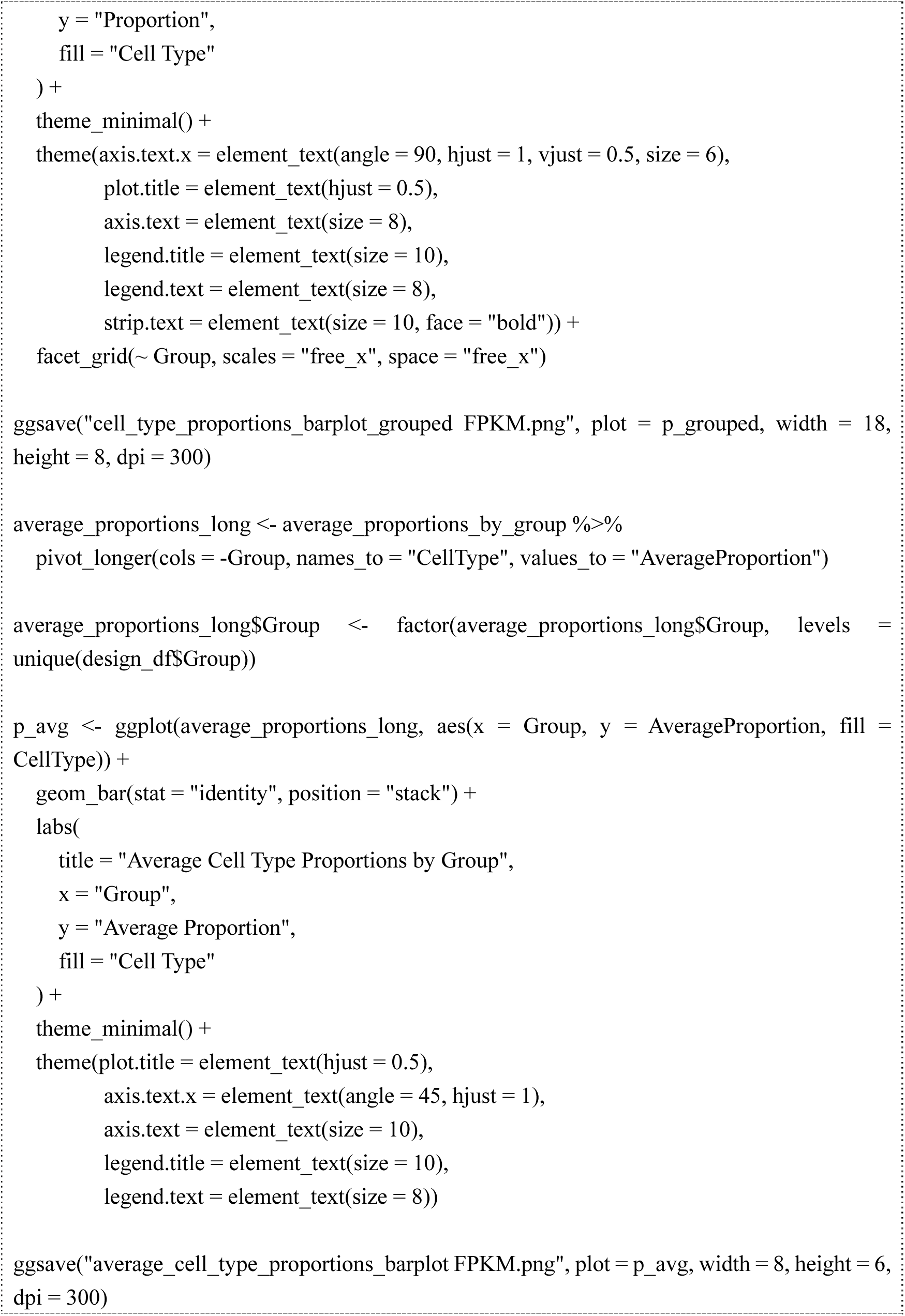

Uncut blots:

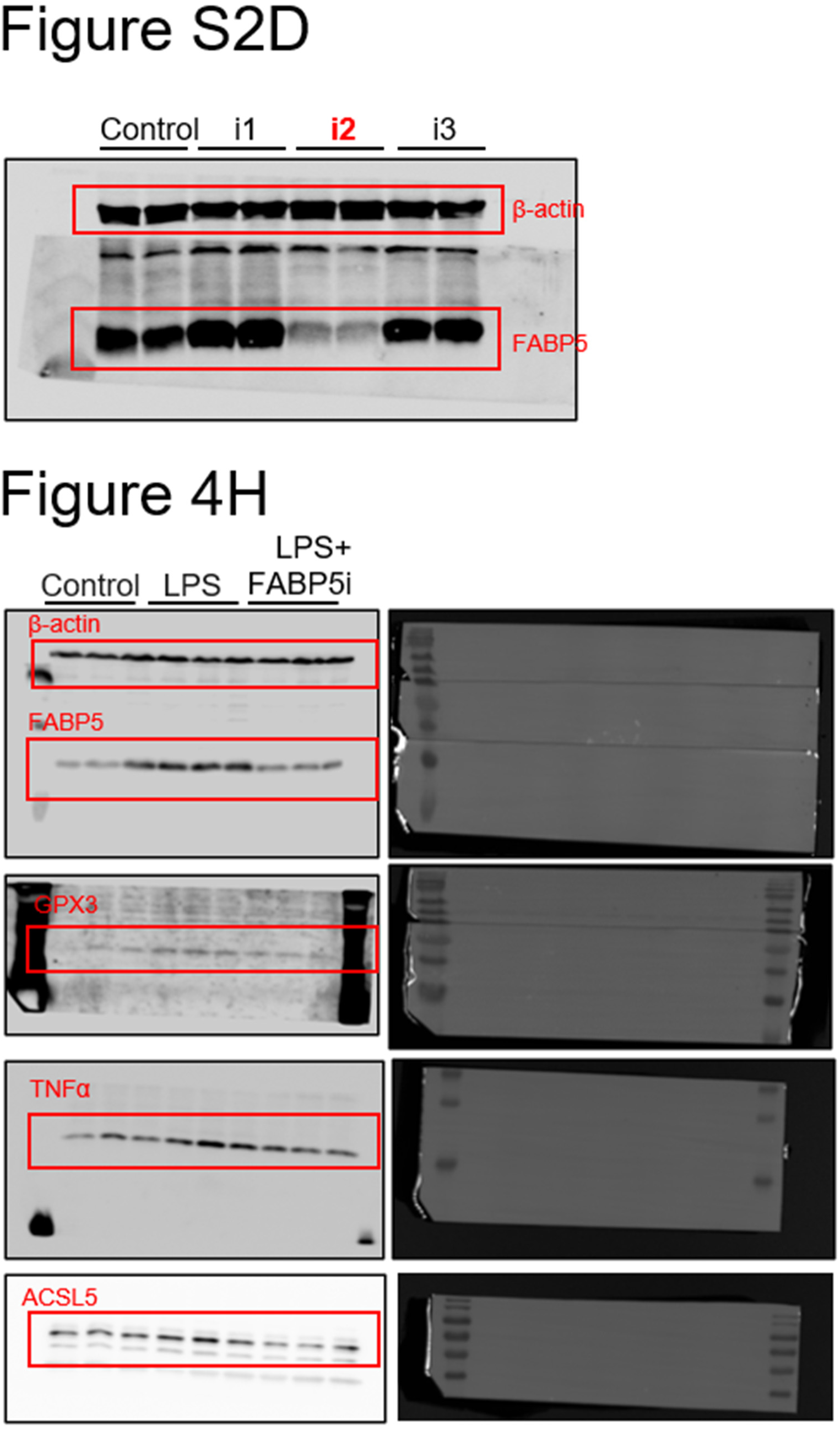

